# The Activome: multiplexed probing of activity of proteolytic enzymes using mass cytometry-compatible activity-based probes (TOF-probes)

**DOI:** 10.1101/775627

**Authors:** Marcin Poreba, Katarzyna Groborz, Wioletta Rut, Milind Pore, Scott J. Snipas, Matej Vizovisek, Boris Turk, Peter Kuhn, Marcin Drag, Guy S. Salvesen

## Abstract

The activome can be considered as a subset of the proteome that contains enzymes in their catalytically active form and can be interrogated by using probes targeted towards individual specific enzymes. A subset of such enzymes are proteases that are frequently studied with activity-based probes, small inhibitors equipped with a detectable tag, commonly a fluorophore. Due to the spectral overlap of these commonly used fluorophores, simultaneous analysis becomes limited. To overcome this, we developed a series of protease-selective lanthanide-labeled probes compatible with mass cytometry. Using lanthanide-based tags instead of fluorophores gives us the ability to monitor the activity of multiple proteases in parallel. As proof of concept we developed a panel of cathepsin and legumain specific probes and showed that we were able to identify an activome of these proteases in two cell lines and peripheral blood mononuclear cells, providing a framework for the use of mass cytometry for multiplexed enzyme activity detection.

## Introduction

Proteases play critical roles in multiple processes in health and disease [1]. Since proteases are characterized by the ability to stimulate peptide bond breakdown, their analysis through genomics or transcriptomics is limited as those do not consider whether the enzyme is active or not [2, 3]. Even proteomic tools such as antibodies that rely on mass spectrometry are often insufficient to indicate which particular protease is active within a proteome [4]. The investigation of protease activity becomes more complicated because these enzymes are regulated on several posttranslational levels [5]. Efforts to explore the universe of proteolytic events must include the identification and activity status of protease(s) in question. Here, we propose a concept called the activome (**Figure 1A**) which can be interrogated by utilizing reagents such as specific inhibitors and activity based probes [2, 6]. The activome holds information about the complete activity status of a sample. Therefore, the activome can be considered as the functional readout of the proteome [2]. The activome does not account for enzymes that were not expressed or are not active due to inhibition, degradation, or lack of proenzyme activation; thus, it extracts information of specific biological relevance. The most convenient approach to study the activome is to use activity-based probes (ABPs) which are small-molecule inhibitors equipped with a detectable tag (biotin, fluorophore, or radioisotope) (**Figure 1B**) [7, 8]. Fluorescently tagged ABPs provided a breakthrough in revealing activomes in cells and whole organisms [9, 10]. However, the overlap of fluorescence emission spectra limits the number of enzymes that can be detected and visualized in parallel [11], thus limiting their application for multi-parametric analysis. To increase the number of enzymes that can be detected simultaneously in the activome we developed metal (lanthanide) isotope-tagged ABPs that can be investigated by both, mass and imaging mass cytometry (IMC). These technologies combine flow cytometry with high precision mass markers, thus overcoming the problem of signal overlap and provide a spatial resolution of 1μm^2^ [12-14]. As elemental mass spectrometry allows for metal isotope discrimination with high precision, it can be applied for simultaneous assessment and quantification of cellular processes at the single cell and protein level [15]. CyTOF (Cytometry by Time-Of-Flight) applications incorporate lanthanide metals to label antibodies, proteins, or DNA/RNA, as well as small chemical probes [16, 17]. Nevertheless, to successfully apply mass cytometry to study the activome, a new type of chemical tools is needed.

**Figure 1.**
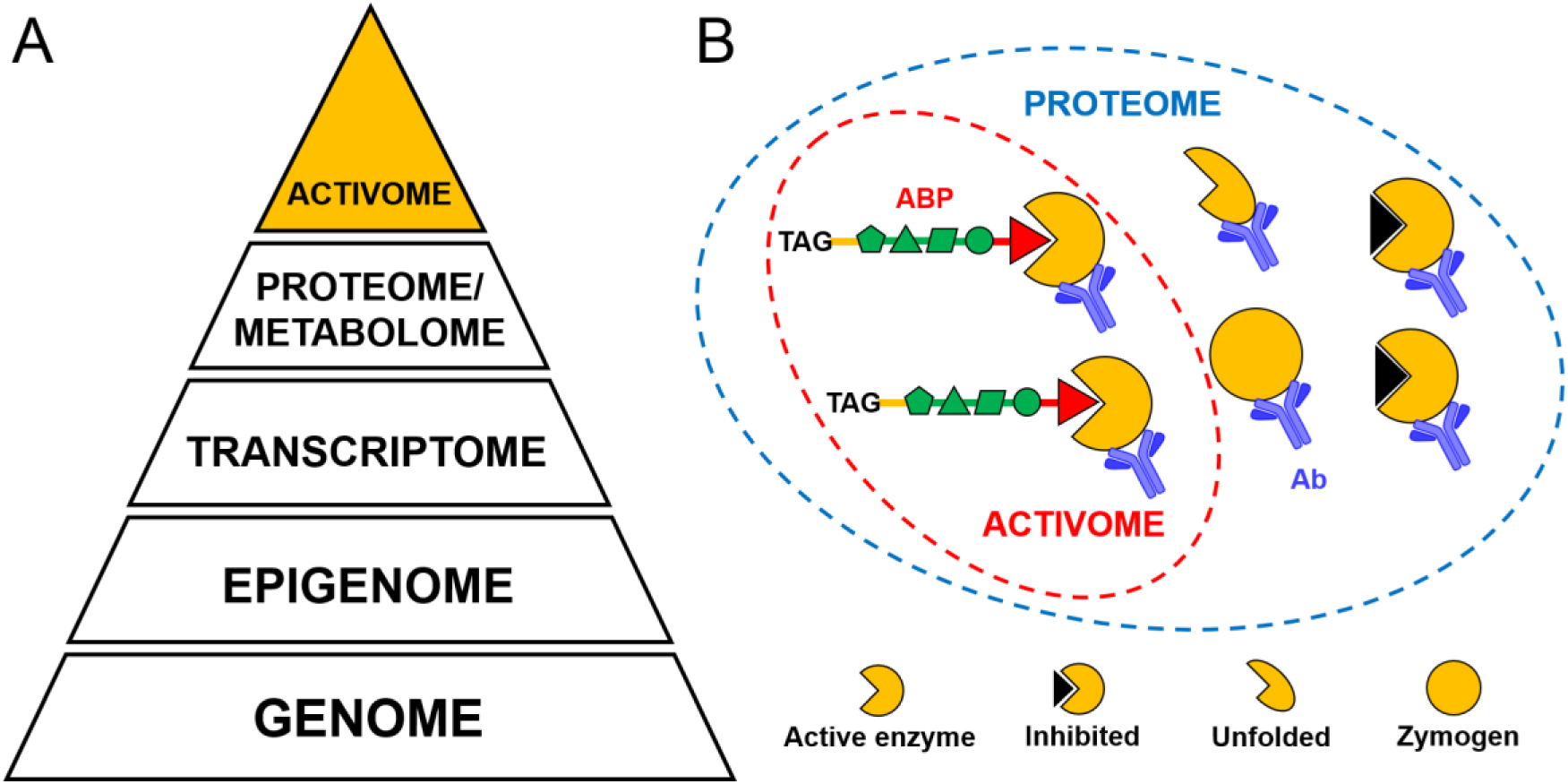
The activome concept. **A** The pyramid represents the hierarchy of experimentally verifiable properties of biological systems, where the apex is defined by the activome: the result of supporting events including translation, expression, and posttranslational modifications. **B** While the proteome is the whole set of proteins regardless of their activity status, the activome is the functional part of the proteome, and can be dissected by selective activity-based probes (ABPs).

The proteases cathepsin B, L and legumain have unique activities and are each involved in protein processing and antigen presentation. Previous studies have defined enzyme-selective sequences that can be utilized to derive metal-mass tagged ABPs [18-20]. Accordingly, we used these three proteases as targets for the development of enzyme-specific TOF-probes (metal-tagged, time of flight activity-based probes, *TOF-probes*). The newly developed probes enabled us to investigate the cellular activity and localization of three lysosomal proteases, cathepsin B, cathepsin L and legumain, using standard mass cytometry and imaging mass cytometry. This work seeks to provide the first example of a mass cytometry platform for parallel enzyme detection, thus setting the stage for future activome research.

## Results

In our previous work we had deployed HyCoSuL to develop highly selective peptide recognition sequences for cathepsin L [18], cathepsin B [21] and legumain [19] (**Figure S1**). Next we aimed to integrate these sequences into a mass cytometry platform by incorporating an *N*-terminal tetracarboxylic acid (DOTA)-chelated stable isotope of lanthanides, and a *C*-terminal acyloxymethylketone (AOMK) warhead (**Figure 2A, B**). TOF-probes were synthesized by coupling a warhead moiety to a DOTA-peptide fragment and chelating corresponding metal particle (**Figure 2C**). To validate our chemical approach, we selected more than one stable metal isotope for each TOF-probe targeting particular enzyme and created a panel of chemical compounds (**Figure 2D**). We decided to use two stable and isotopically pure metals-terbium and lutetium (^159^Tb and ^175^Lu), and naturally occurring gadolinium, which is a mixture of six stable isotopes (^154^Gd, ^155^Gd, ^156^Gd, ^157^Gd, ^158^Gd, ^160^Gd). The mixture of gadolinium isotopes was used in order to evaluate whether the nature of the isotope can influence enzyme binding specificity.

**Figure 2.**
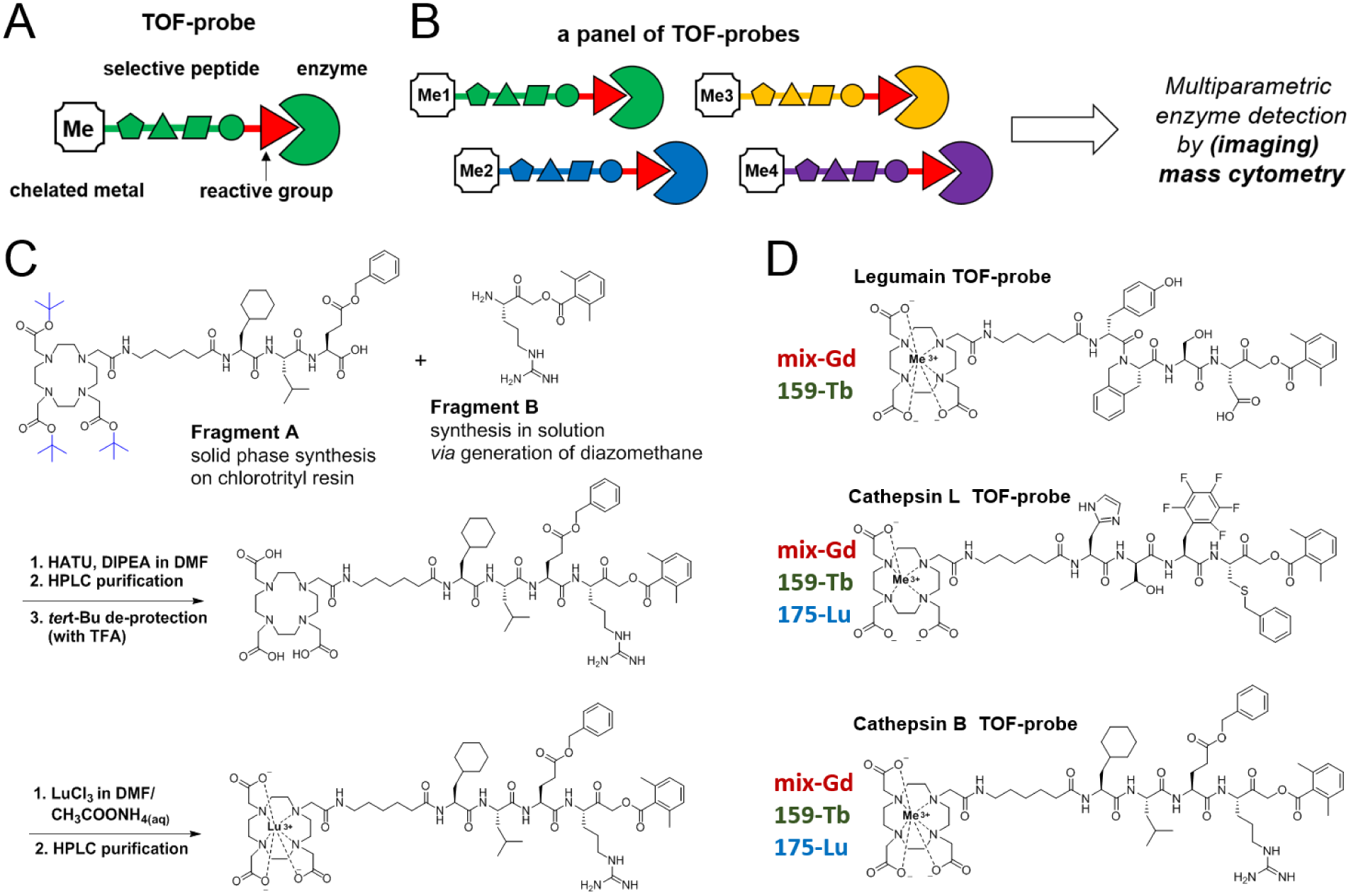
The concept of TOF-probes for the selective detection of enzymes with mass cytometry. **A** The general architecture of a TOF-probe. A protease-selective peptide is tagged with a stable lanthanide isotope making it suitable for mass cytometry. **B** A set of protease-selective probes labeled with different metal tags for the parallel visualization of several enzymes. **C** An outline of the chemical procedure for the high yield synthesis of TOF-probes. **D** Examples of protease-selective TOF-probes labeled with different lanthanides. In this work we synthesized a panel of eight TOF-probes for three proteases (legumain, cathepsin L and cathepsin B) based on previously developed selective peptide sequences.

Fluorescent ABPs are valuable tools for the detection of enzyme activity, however, one of their main limitations is the fluorescent spectral overlap, which largely reduces the number of enzymes that can be visualized simultaneously (**Figure S2**) [11]. In a quest to develop a new method for multiplexed enzyme imaging, we designed ABPs that are compatible with mass cytometry, a method that uses stable isotopes of lanthanide metals as chemical tags [12]. DOTA-metal tagged TOF-probes provided good activity and specificity for the individual enzymes (**Figure 3**). Moreover, regardless of the metal tag used (Lu, Gd, Tb), TOF-probes retained their binding potency (**Figure 3A**). This strongly suggests that all available metals can be interchangeably used for TOF-probe labeling, making them compatible with commercially available metal-tagged antibodies. We determined the utility of TOF-probes in cells by competition against fluorescent probes. Gd-tagged TOF-probes for legumain, cathepsin L or cathepsin B were incubated at various concentrations in either MDA-MB-231 or HCT-116 cell lines, followed by measuring the residual activity of selected enzymes (**Figure 3B, C**; **Figure S3**). This experiment revealed that 2 μM legumain TOF-probe was sufficient to selectively block legumain activity in both cell lines. Cathepsin L and B TOF-probes were also highly potent and although we observed cross-reactivity between them at high concentrations, at 2 μM probes were highly selective for their target enzymes.

**Figure 3.**
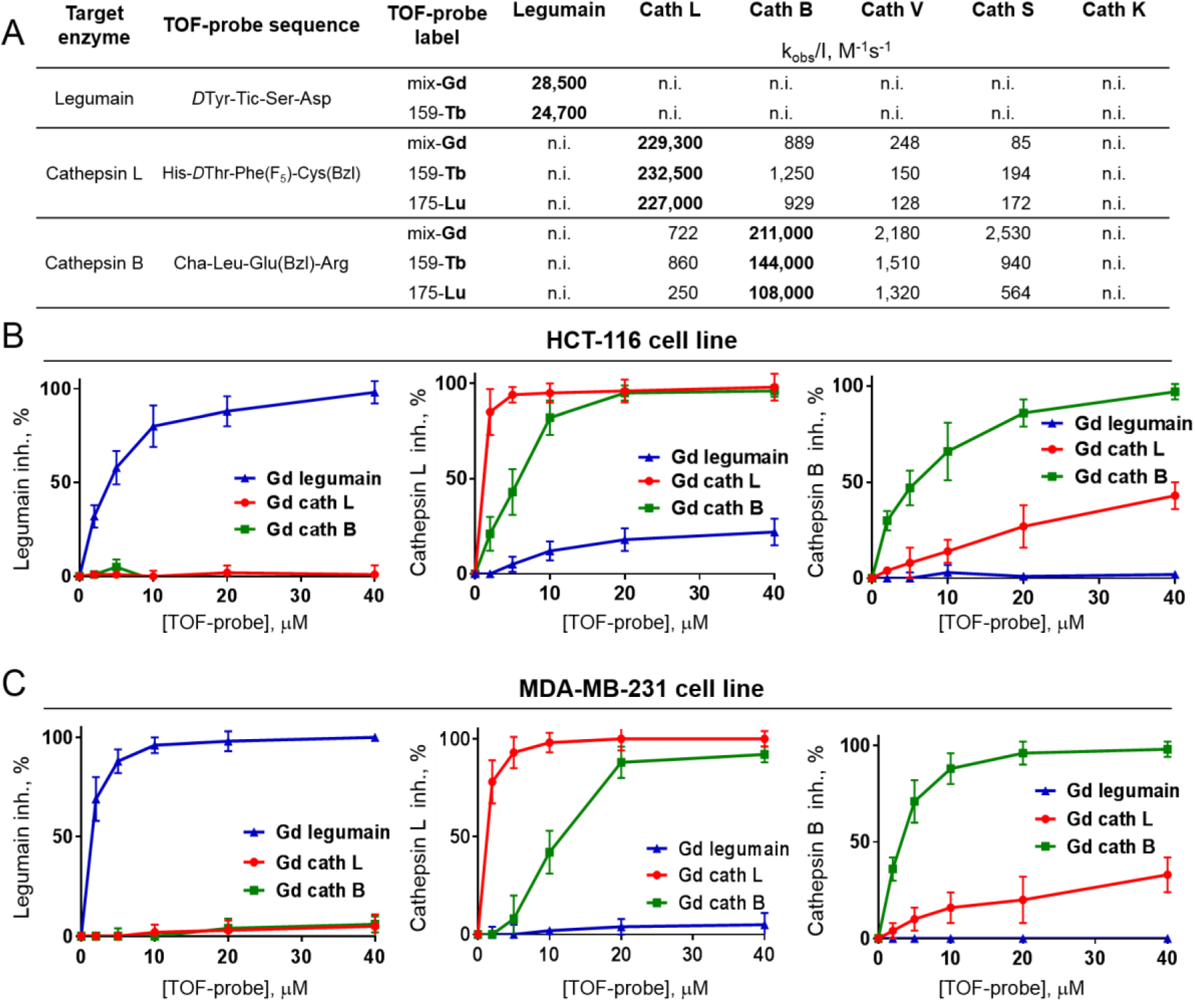
Specificity of TOF-probes toward recombinant proteases and cancer cell lines. **A** Kinetic parameters (k_obs_/[I]) of eight TOF-probes measured against three human recombinant proteases (legumain, cathepsin L and cathepsin B). **B** and **C** Selectivity of TOF-probes toward target proteases in cancer cell lines HCT-116 (B) and MDA-MB-231 (C). To assess the selectivity of TOF-probes, they were incubated at various concentration ranges with two cancer cell lines, and residual protease activity was detected with selective, Cy5-labeled probe (Cy5-MP-L01 for legumain, Cy5-MP-cL3 for cathepsin L and Cy5-cB2 for cathepsin B). Based on signal intensity, TOF-probe inhibition profiles were determined. The data demonstrated that regardless of metal tag, all TOF-probes are selective toward targeted proteases.

Having confirmed the potency, selectivity and cellular uptake of TOF-probes, we employed these probes to explore the activome by using mass cytometry. After incubation with probes, cells were washed and fixed to remove excess probe. We observed the individual signals from simultaneously labelled samples (**Figure 4**). Specificity of the TOF-probes in this experiment was confirmed by competition with inhibitors, demonstrating>10-fold signal-to-noise ratio (**Figure 4C**). All cells were stained with Ir191/Ir193 DNA intercalator to detect cell events. By using the cathepsin B-directed, Gd-labeled-TOF-probe, which can be detected in several channels, we demonstrated that the individual probe is evenly distributed across the cells (**Figure S4**). We found that the efficiency of cell labeling is time-dependent, as after 1 hour of incubation around 20% of cells were probe positive, increasing to over 80% after an additional 3 hours (**Figure S4).**

**Figure 4.**
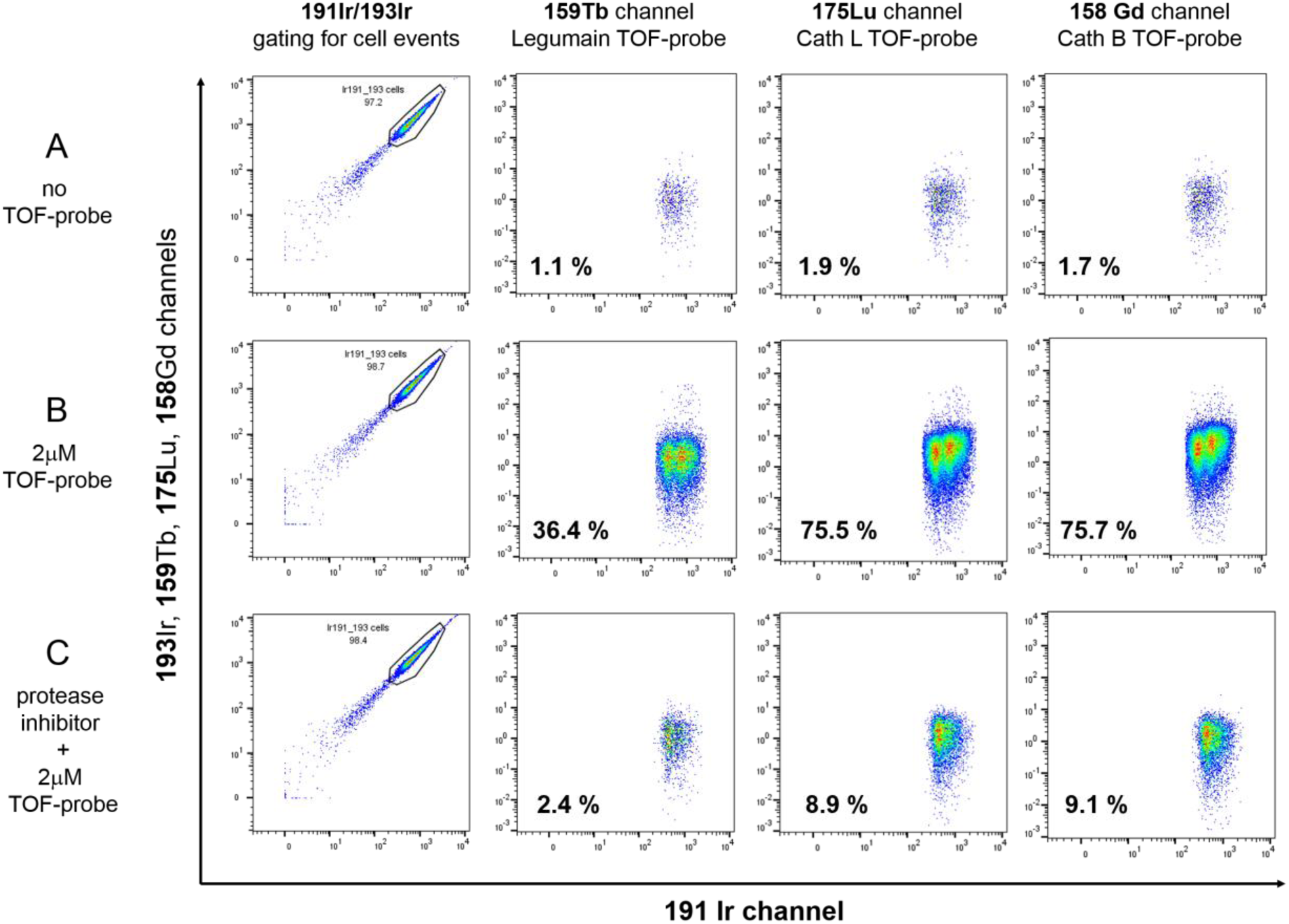
Detection of proteases by TOF-probes in HCT-116 cells. TOF-probes that were incubated with cells are taken up and react covalently with proteases. After fixing and permeabilization, unbound probes are washed out. For each measurement around 100,000 cell events were collected and analyzed. Cells were (**A**) left untreated to measure background signal, (**B**) incubated with TOF-probes (2 μM, 6 hours), or (**C**) pre-treated with inhibitors (25 μM, 2 hours), followed by incubation with TOF-probes (2 μM, 6 hours). Cells were stained with ^191^Ir/^193^Ir (DNA intercalator to reveal the nucleus) and subjected for mass cytometry analysis. Cells that were left untreated have a low background signal, whereas cells with TOF-probes demonstrate labeling of proteases: legumain (^159^Tb channel), cathepsin L (^175^Lu channel) and cathepsin B (^158^Gd channel). When the cells were pre-treated with inhibitors (MP-L01 for legumain and E64d for cathepsins) signals from enzymes were substantially reduced. For each sample the percentage of cell positive events was calculated.

To verify that metals in the TOF-probes can be used interchangeably, we stained HCT-116 cells with three combinations of TOF-probes (Gd, Tb, and Lu) (**Figure 5)**. The results revealed that the pattern of protease labeling was similar across all TOF-probe variants (**Figure 5**). These data validate the principle of using TOF-probes to investigate subsets of the activome in parallel (in this case cathepsin B, L and legumain). Given the fact that the metal tags had no impact on the cellular uptake of TOF-probes and the corresponding protease detection, our approach is highly compatible with other mass cytometry labeling reagents. Detailed gating strategy is shown in **Figure S5** and **Figure S6**.

**Figure 5.**
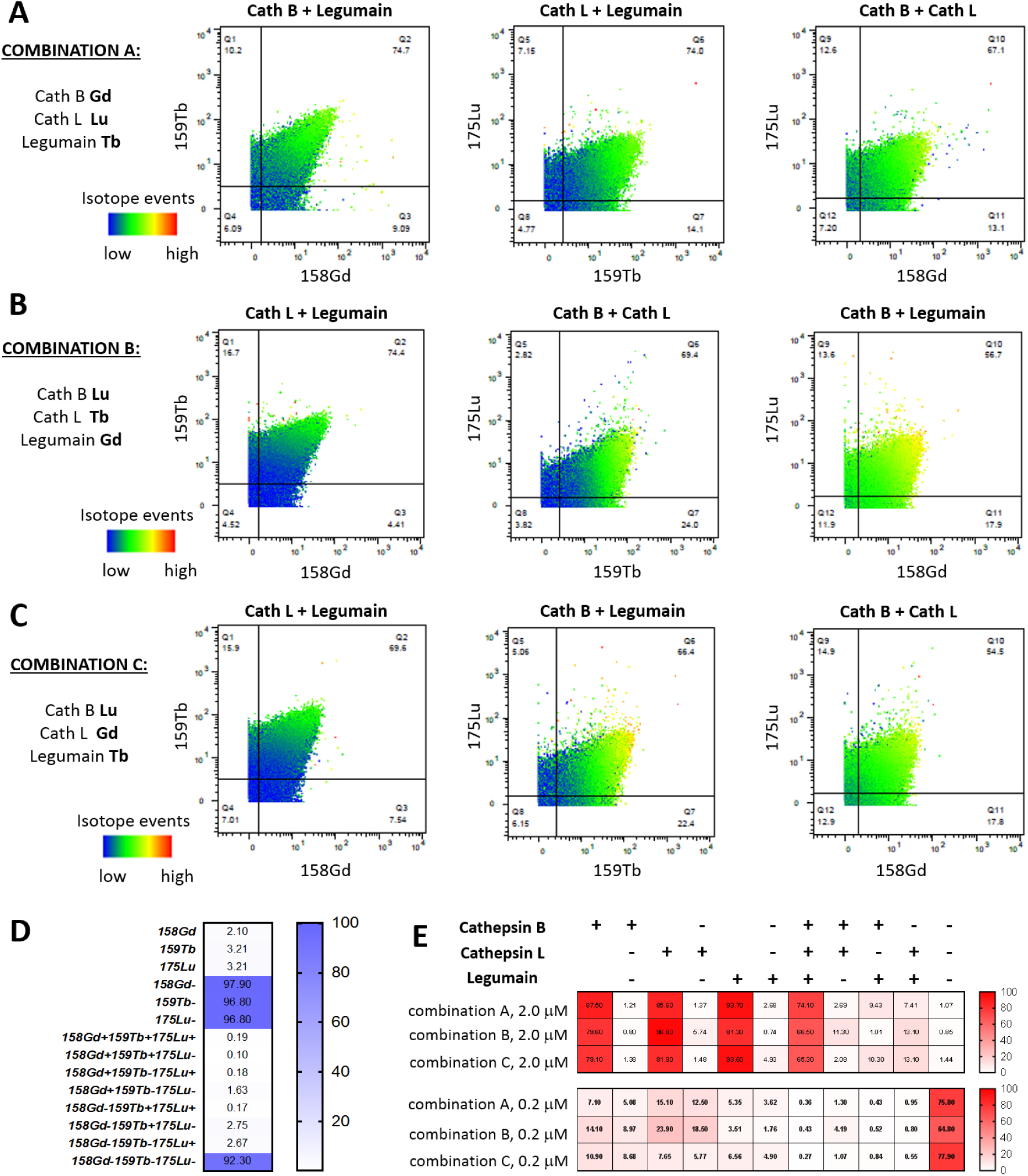
Selective labeling of proteases in HCT-116 cells with TOF-probes. **A-C** Cells were incubated with various sets of TOF-probes (in three combinations: A-C). Graphs were divided into four quadrants based on metal (TOF-probe) events. **D** Non-TOF-probe treated cells were subjected to mass cytometry analysis and the data was used for TOF-probe treated cell gating. **E** TOF-probes uptake and protease labeling in HCT-116 cells presented as heat maps. Data demonstrate that regardless of which TOF-probes set was used, the pattern of protease labeling is similar, demonstrating the wide applicability of this technology.

THP-1, a non-adherent monocyte-like cell line, a model of mononuclear cells, is predicted to transcribe low levels of legumain and therefore serves as a good platform for determining the sensitivity of TOF-probes. Using the same conditions as in previous experiments (**Figure 4, Figure 5, Figure S7**) revealed that cathepsin B and L were abundant (**Figure 6**) mirroring the transcription data shown in **Figure S8**. On the other hand, the staining of legumain was poor (only 15.6%), which reflects the low expression of this enzyme (**Figure 6, Figure S8**). Moreover, pre-incubation of THP-1 cells with inhibitors substantially reduced the signals, demonstrating that TOF-probes displayed very little off-target labeling or non-selective accumulation in the lysosomal compartments (**Figure 6B,C**).

**Figure 6.**
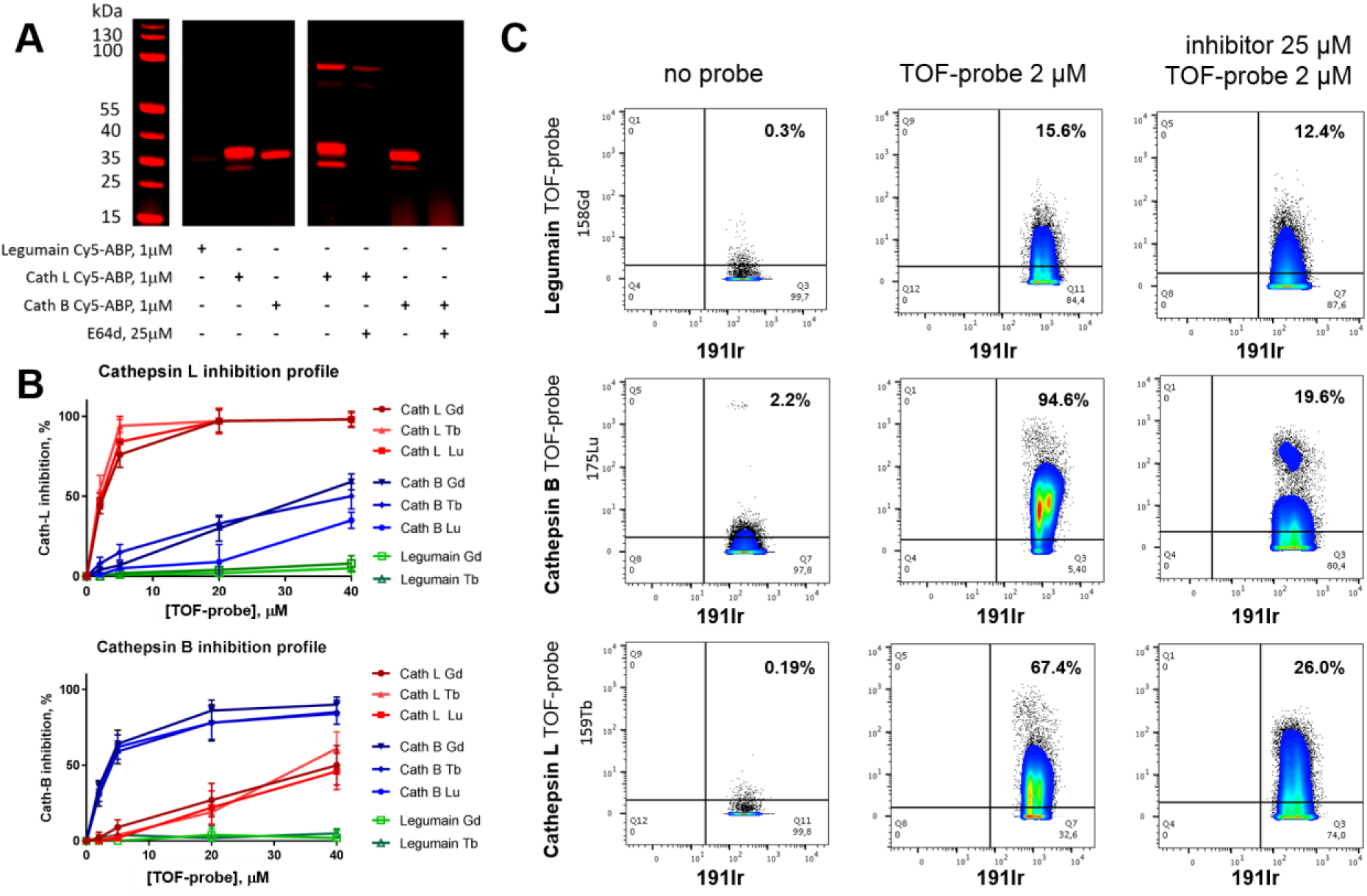
Detection of protease activity in THP-1 cells using TOF-probes**. A** Visualization of cathepsin L and cathepsin B activity in THP-1 with Cy5-labeled selective probes. Legumain was not labeled with Cy5-MP-L01 probe, confirming the low expression level of this enzyme in THP-1 cells. **B** Inhibition profiles of cathepsin L and cathepsin B in THP-1 cells. Data demonstrate high selectivity of our TOF-probes. **C** Metal-labeled TOF-probes show high activity of cathepsin L and cathepsin B, but not legumain in THP-1 cells. Left panel: background signal from investigated channels; middle panel: the percentage of cells containing active proteases; right panel: residual protease activity after inhibitor pre-treatment.

Having demonstrated the applicability of TOF-probes in identifying cellular populations we wondered whether this technology would be able to reveal subcellular structures of the cathepsin and legumain activome. Accordingly, we imaged TOF-probe treated THP-1 cells by high-resolution laser ablation coupled to mass cytometry (IMC) (**Figure 7A**) [22]. Since the cells are dried during sample preparation, membrane and cytoplasmic components coincide with the nucleus, which defines the segmented area. The images reveal a punctate staining pattern for each of the probes consistent with a lysosomal location for the target proteases, as observed previously using fluorescent ABPs [18-20]. This interpretation is verified by IMC experiments previously performed with antibodies coupled to lysosomal membrane proteins (LAMP-2) [23]. The probe enrichment observed in the nuclear region is likely due to cell dehydration during sample preparation. Thus, imaging mass cytometry is a valuable technology for revealing the spatial distribution of the cellular activome (**Figure 7B**).

**Figure 7.**
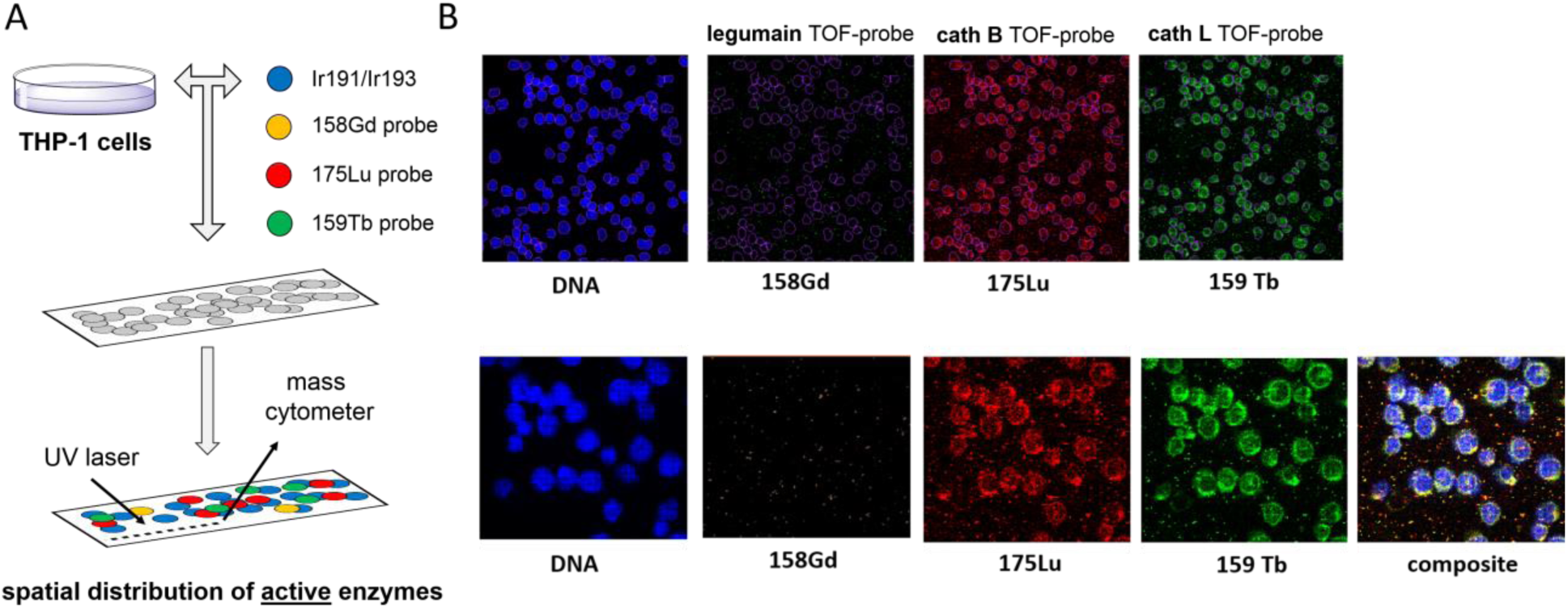
Spatial distribution of active proteases in THP-1 cells. **A** THP-1 cells were attached to slides and incubated with a panel of TOF-probes. After TOF-probe treatment, cells were fixed, permeabilized and incubated with ^191^Ir/^193^Ir to label DNA. Next, slides were subjected for laser ablation with imaging mass cytometry (IMC) to reveal the spatial distribution of active proteases. **B** Detection of active cathepsin B and cathepsin L in THP-1 cells with TOF-probes. No active legumain was detected.

To develop the utility and demonstrate the application of our approach we explored the cathepsin B, L and legumain activome in peripheral blood mononuclear cells (PBMC). PBMCs were incubated with TOF-probes, and metal-tagged cell surface receptor antibodies to detect individual cell populations (T cells, B cells, natural killer (NK) cells) (**Figure 8)**. The data showed that T, B, and NK cells contain equivalent amounts of active cathepsins B and L. Active legumain is abundant in B cells compared to the other cell types. The data are consistent across different PBMC preparations and not dependent on the metal used in the TOF-probe structure (**Figure 8A, Figure S9, S10**). To test whether TOF-probes were being taken up non-specifically we increased the concentration of each to 10 μM (**Figure 8B)**. Although the portion of cells labelled with cathepsin B and L probes increased substantially, the portion of cells labelled with a legumain probe increased by only 5.4% in T cells. We interpret this to signify that most PBMCs contain relatively large amounts of cathepsin B and L, but small amounts of legumain, and that our sample washing protocol is sufficient to remove excess probe. Cathepsin L and cathepsin B TOF-probes enabled us to divide the NK cells into two distinct populations: positive, and high positive, demonstrating that the protease activome may vary across the same type of cells as previously seen with neutrophils [11]. The labeling of PMBCs with 10 μM legumain TOF-probe confirmed that this protease is active in B cells (77.9% positive), moderately active in NK cells (39.9%), but poorly active in T cells (19.1%). Whether this differential is of biological significance is unclear, but merits further investigation.

**Figure 8.**
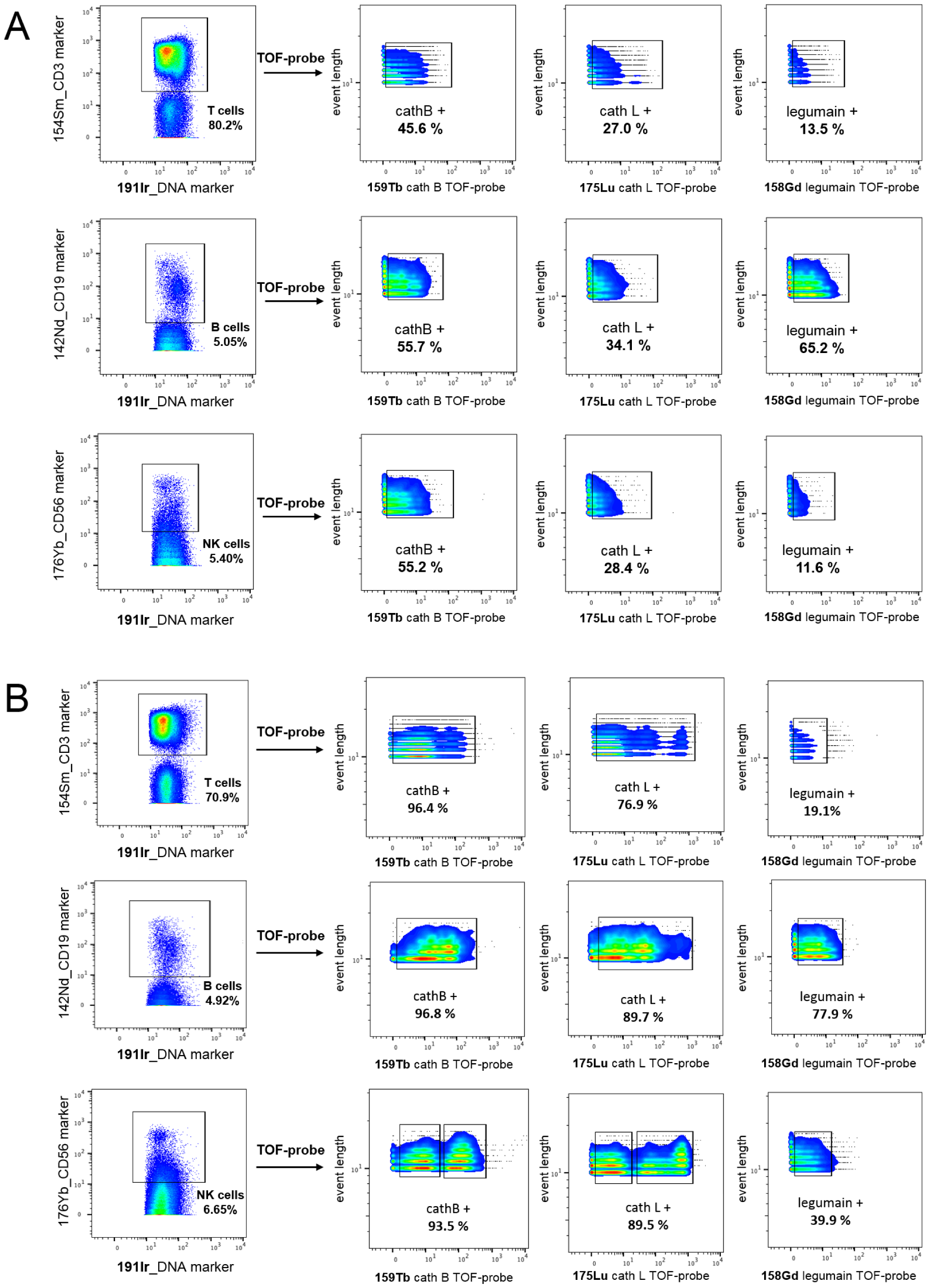
Protease detection in Peripheral Blood Mononuclear Cells (PBMCs). **A** Active proteases were labeled with TOF-probes (2 μM) in PBMCs. The data show that cathepsin B is the most active enzyme in all type of cells, cathepsin L is also present in all cell populations, but less active than cathepsin B, and legumain activity is B cells specific. **B** Active proteases were labeled with TOF-probes (10 μM) in PBMCs. The results show that cathepsin B and cathepsin L were highly active in all types of cells, whereas legumain was mainly found in B cells, and to a lesser extent in NK cells.

## Discussion

The development of selective probes to dissect the activity of individual proteases within a complex system from the test-tube all the way to *in vivo* imaging in whole animals has witnessed substantial progress over the last 20 years [9, 24, 25]. The development of increasingly selective probes has enabled the tracking of multiple proteases that together form an activome, a functional component of the proteome. However, one of the major remaining limitations for the parallel monitoring of individual proteases is the lack of appropriate tagging strategies. Currently, most ABPs contain fluorophores that display narrow excitation/emission spectra and good quantum yield are mostly located within a relative narrow spectral range between 500 – 800 nm. Therefore, these fluorophores allow for the visualization of only up to 4-5 proteases in parallel to avoid substantial spectral overlap [11]. We reasoned that employing Lanthanide-based MS-tags for ABP labeling would enable us to study multiple proteases at the same time and expand our ability to interrogate cellular activomes. In this study, we developed metal-tagged ABPs for three proteases, for which selective probe-targeting peptide sequences had been already reported [18, 19]. Our analysis revealed that these new probes, which we called TOF-probes, display high potency and selectivity towards their enzyme targets. To validate their utility, we used them for simultaneous detection of individual proteases in three cell lines using mass cytometry. This experiment demonstrated that our TOF-probes are efficiently taken up by the cells, and selectively label proteases of interest. We determined that the metal tags can be used interchangeably, as the nature of the tag does not affect the TOF-probes potency and selectivity. This makes the approach compatible with metal-tagged antibodies or DNA probes that are currently used for tagging cells for mass cytometry. One difference between TOF-probes and metal-tagged antibodies is that former are able to chelate only one metal particle per probe, whereas antibodies labelled with lanthanides contain up to dozen metal particles attached to the polymer. Therefore, metal-labelled antibodies generate significantly higher signals that may provide for more evident interpretation. Nevertheless, by introducing apropriate controls we are able to dissect positive signals in the samples treated with TOF-probes. Going one step further, we also detected protease activity in cells using imaging mass cytometry (IMC), a technique that enables the simultaneous, multi-parametric analysis of a sample. We showed that active cathepsin B and cathepsin L strongly overlap in THP-1 cells, whereas active legumain is very low to undetectable in these cells. In this experiment, we also demonstrated that TOF-probes can be integrated with IMC settings to create a chemical platform to resolve the spatial distribution of multiple active enzymes within cells or tissue microenvironments.

To demonstrate the broad applicability of the newly developed platform, we aimed to decipher the cathepsins and legumain activome fingerprint in individual populations. Metal-tagged cell antibody markers were used to distinguish individual leukocyte populations (T cells, B cells, and NK cells) in combination with our TOF-probes. This analysis showed that cathepsins B and L are abundantly active in resting leukocytes, whereas the activity of legumain is cell type dependent, being the most active in B cells. Knowledge of the activity of individual proteases, or enzymes in general, in specific cell populations has the potential to translate into therapeutic, theragnostic, or biomarker settings. For example, the analysis of legumain activity in overall PBMC populations is misleading, as T-cells are the most abundant cells in PBMC population and they display only negligible legumain activity. However, by combining TOF-probes with cell markers it becomes possible to integrate proteolytic activity across heterogenous cell populations to generate a more holistic understanding of the activome and more informative protease readouts. With the increased number of deployable metal isotopes it will be possible to gain insight into the complexity of enzyme activities within healthy tissues and disease lesions at a single cell level to provide system-wide views of their activation and their role in complex biological networks.

## Material and methods

### Chemical reagents

All chemicals used for the synthesis of metal-labeled activity-based probes (TOF-probes) were purchased from commercial suppliers and used without purification unless otherwise noted. The 2-cholotorityl chloride resin (1.59 mmol/g, 100-200 mesh) was used for the synthesis of peptides that were further converted into TOF-probes (Iris Biotech GmbH, Germany). Fmoc-protected amino acids were purchased from various suppliers: Iris Biotech GmbH, P3 BioSystems (Louisville, USA), QM Bio (Shanghai, China), and Bachem (Bubendorf, Switzerland). Diisopropylcarbodiimide (DICI, peptide grade), N,N-diisopropylethylamine (DIPEA, peptide grade), piperidine (PIP, peptide grade), and trifluoroacetic acid (TFA, purity 99%) were all from Iris Biotech. 2,4,6-trimethylpyridine (2,4,6-collidine, peptide grade), triisopropylsilane (TIPS, purity 99%), 2,2,2-trifluoroethanol (TFE), anhydrous tetrahydrofuran (THF), hydrogen bromide (30% wt. in AcOH), 4-methylmorpholine (NMM), isobutylchloroformate (IBCF), and 2,6-dimethylbenzoic acid (2,6-DMBA) were all purchased from Sigma Aldrich. N-hydroxybenzotriazole (HOBt, monohydrate) was from Creosalus (Louisville, USA). HATU and HBTU (both peptide grade) were from ChemPep Inc.. N,N’-dimethylformamide (DMF, peptide grade) and acetonitrile (ACN, HPLC pure) were from WITKO (Lodz, Poland). Methanol (MeOH, pure for analysis), dichloromethane (DCM, pure for analysis), diethyl ether (Et_2_O, pure for analysis), acetic acid (AcOH, 98% pure) and phosphorus pentoxide (P_2_O_5_, 98% pure) were from POCh (Gliwice, Poland). Fluorescent tags (Cyanine-5 NHS and Cyanine-7 NHS) were purchased from Lumiprobe (Hannover, Germany). Diazomethane was generated according to the Aldrich Technical Bulletin (AL-180) protocol.

### Synthesis of fluorescent activity-based probes

The detailed protocol for the synthesis of Cy5-labeled ABPs for legumain, cathepsin L and cathepsin B is published elsewhere [18]. All synthesized ABPs were purified on HPLC, and their MS was confirmed via HR-MS.

### Synthesis of TOF-probes (metal-tagged protease inhibitors)

All the amino acids presented in this procedure are *L*-enantiomers, unless otherwise stated. The detailed procedure of TOF-probes synthesis is presented on the example of DOTA(^175^Lu)-Ahx-Cha-Leu-Glu(Bzl)-Arg-AOMK (for cathepsin B, Figure 2C), and other TOF-probes utilized in this work were obtained in the same way. The synthesis of ^175^Lu-tagged cathepsin B probe included five sequential stages: (1) Boc-Arg(Boc)_2_-AOMK warhead synthesis, (2) the synthesis of selective peptide sequence with 6-aminohexanoic acid spacer (Boc-Ahx-Cha-Leu-Glu(Bzl)-OH); (3) coupling of warhead with peptide sequence and de-protection of amino acids side chains (H2N-Ahx-Cha-Leu-Glu(Bzl)-Arg-AOMK); (4) conjugation of DOTA chelating agent to obtain DOTA-Ahx-Cha-Leu-Glu(Bzl)-Arg-AOMK; (5) and incorporation of trivalent metal isotope (DOTA(^175^Lu)-Ahx-Cha-Leu-Glu(Bzl)-Arg-AOMK). The irreversible reactive group Boc-Arg(Boc)_2_-AOMK was obtained according to the procedure described previously [19, 26]. Cathepsin B selective peptide sequence was synthesized on the solid support (2-chlorotrityl chloride resin) according to standard solid phase peptide synthesis method [27]. In brief, Fmoc-Glu(Bzl)-OH (3 eq) was coupled to the resin (1 eq, 250 mg) in anhydrous DCM using DIPEA (3 eq) within 3 h. Then, Fmoc-protecting group was removed with 20% piperidine in DMF (5, 5, 25 min cycles). In the next steps Fmoc-Leu-OH, Fmoc-Cha-OH and Boc-6-Ahx-OH (2.5 eq) were attached using HATU (2.5 eq) and 2,4,6-collidine (2.5 eq) in DMF as coupling reagents. Each amino acid coupling was carried out for 2.5 h. In the last step, the peptide was removed from the resin with the mixture of DCM/TFE/AcOH (v/v/v, 8:1:1) within 45 min. Then, the solution was filtered and solvents were removed under reduced pressure. The crude peptide was dissolved in acetonitrile:H_2_O (v/v, 3:1) and lyophilized to obtain Boc-6-Ahx-Cha-Leu-Glu(Bzl)-OH as a white powder (purity ≥ 95%). In the next stage H_2_N-Arg-AOMK was coupled to Boc-6-Ahx-Cha-Leu-Glu(Bzl)-OH. To a small flask containing 1 eq of Boc-Arg(Boc)_2_-AOMK the mixture of DCM:TFA (v/v, 2:1) was added to remove protecting groups. After 30 min TFA and DCM were evaporated under reduced pressure. To freshly obtained H_2_N-Arg-AOMK (yellow oil) Boc-6-Ahx-Cha-Leu-Glu(Bzl)-OH (1.2 eq), HATU (1.2 eq) in DMF and 2,4,6-collidine (3 eq) were added. The coupling reaction was monitored by HPLC. After the reaction was completed (3 h) the mixture was diluted with ethyl acetate, transferred to a separatory funnel and extracted with 5% citric acid (once), 5% NaHCO_3_ (once) and brine (once). The organic fraction was dried over MgSO_4_ and evaporated under reduced pressure. Obtained Boc-Ahx-Cha-Leu-Glu(Bzl)-Arg-AOMK was treated with the mixture of DCM:TFA (1:1, v/v) for 30 min and the solvents were removed under reduced pressure. Crude product was purified on HPLC and lyophilized (purity ≥ 95%). In the next stage, DOTA(*t*Bu)_3_ (1.2 eq) and HATU (1.2 eq) were dissolved in a small volume of DMF. Then, collidine (3 eq) was added to this solution. The mixture was activated for 1 min and added to a small flask containing 1eq of H2N-6-Ahx-Cha-Leu-Glu(Bzl)-Arg-AOMK in DMF. The reaction mixture was agitated at room temperature until HPLC indicated that the reaction was complete (one hour). Then, obtained DOTA(*t*Bu)_3_-Ahx-Cha-Leu-Glu(Bzl)-Arg-AOMK was purified on HPLC and lyophilized (purity ≥ 95%). In the last stage, the conjugation of DOTA-peptide with metal isotope was carried out according to the method described by Sasabowski and Mather [28]. To remove *t*Bu-protecting groups DOTA(*t*Bu)_3_-Ahx-Cha-Leu-Glu(Bzl)-Arg-AOMK (1 eq) was added to a solution of 50% TFA in DCM and stirred for 30 min. After this time, solvents were removed under reduce pressure. Lutetium (III) chloride (5 eq) was dissolved in minimal volume of 0.1 M ammonium acetate buffer (pH= 5.0) and added to a small flask containing DOTA-Ahx-Cha-Leu-Glu(Bzl)-Arg-AOMK dissolved in minimal volume of DMF. The pH of metal conjugation was in the range of 4 to 6 to avoid reducing the rate of metal complexation (pH<4) or formation of insoluble metal hydroxides (pH>6) [4]. The reaction flask was placed in water bath (55°C) and stirred for 1 h (radiolabeling was monitored by HPLC). Subsequently, the crude product was purified on HPLC and lyophilized (purity ≥ 95%).

### Cysteine cathepsins and legumain

Human recombinant cathepsins B, L, V, S, and K were expressed and purified as published previously [29, 30]. In prior to kinetic studies all cathepsins were active site titrated using E64 inhibitor (Peptide Institute, Japan). Human recombinant pro-legumain (asparaginyl endopeptidase) was purchased from R&D Systems (2199-CY), activated per manufacturer instruction and active site titrated using MP-L01 inhibitor as described by Poreba *et al*. [19].

### Enzymatic kinetic studies

Inhibitors kinetic experiments were performed using CLARIOStar (BMG LABTECH) plate reader operating in fluorescence kinetic mode using 96-well plates (Corning®, Costar®). AMC-labeled fluorescent substrates were screened at 360 nm (excitation) and 460 nm (emission) wavelengths (gain 650). Cathepsins substrate (Z-FR-AMC) was from R&D Systems and legumain substrate (Z-AAN-AMC) was from Bachem. The cathepsins assay buffer composition was: 100 mM sodium acetate, 100 mM sodium chloride, 10 mM DTT, 1 mM EDTA, pH 5.5. Legumain assay buffer was: 50 mM MES, 250 mM NaCl, and pH 5.0. Buffers was prepared at ambient temperature and enzymes kinetic studies were performed at 37 °C. For all probes (fluorescent and TOF-probes) the second rate inhibition constant (k_obs_/I expressed in M^-1^s^-1^) towards human cathepsins B, K, L, S, and V and legumain was measured under pseudo-first order kinetic conditions. Individual proteases were mixed with various concentrations of inhibitors (at least 5-fold excess over enzymes) and 100 μM AMC fluorescent substrate. Experiments were repeated at least three times, and results are presented as an average (S.D. were below 20%).

### Cell culture

In this study we used low passage cell lines HCT-116, MDA-MB-231, and THP-1 purchased from ATCC. HCT-116 cells were cultured in McCoy’s 5A medium, MDA-MB-231 cells were cultured in Dulbecco’s Modification of Eagle’s Medium (DMEM) medium, and THP-1 cells were cultured in RPMI-1640 medium. Each medium was supplemented with 10% of Fetal Bovine Serum (FBS), 2 mM L-glutamine, 100 units/mL penicillin and 100 μg/mL streptomycin. Adherent cells (HCT-116 and MDA-MB-231) were detached from culture plate using trypsin-EDTA (0.25%) – phenol red solution.

### Protease labeling in cancer cells using fluorescent ABPs

50,000 of HCT-116 or MDA-MB-231 cells were seeded into 12-well plates and allow to attach overnight. The next day, 1 μM Cy5-labeled probes for either legumain (Cy5-MP-L01), cathepsin L (MP-cL3) or cathepsin B (MP-cB2) was added to the cells and incubated for 4 hours. Next, cells were harvested, centrifuged at 500 × g for 5 min, cell pellets were washed with 1xDPBS, cells were centrifuged again at 500 × g for 5 min, and the supernatant was discarded. Finally, cell pellets were solubilized into 100 μL of 1x SDS/DTT and boiled for 5 min, cooled to room temperature and sonicated. Subsequently, each sample (30 μL) was subjected for SDS-PAGE analysis (4-12% Bis-Tris Plus 10-well gels, 200 V, 30 min; 3 μL of pre-stained protein ladder was used in the first lane). Gels were scanned at 700 nm (red channel for Cy5 detection) using the Odyssey fluorescence imaging system (LI-COR). Images were analyzed in Image Studio software. This verified that each Cy5 probe specifically labeled its target protease. In a similar manner, we labeled proteases in THP-1 cells. In brief, 100,000 of non-adherent THP-1 cells were placed into 12-well plates, followed by incubation with 1 μM Cy5 probes (4 hours). After this time cells were subjected for SDS-PAGE analysis, and gels were scanned as described above.

### Assessment of TOF-probes selectivity and potency in cancer cells

100,000 of HCT-116 or MDA-MB-231 cells were seeded into 12-well plates and allowed to attach overnight. The next day, selected TOF-probes were incubated with cells at five different concentrations (0, 2, 5, 10, 20, and 40 μM) for 4 hours. After this time, 1 μM Cy5-labeled probe was added and incubated with cells for additional four hours to label residual protease activity (Cy5-MP-L01 for legumain, MP-cL3 for cathepsin L and MP-cB2 for cathepsin B). Next, cells were harvested and subjected to SDS-PAGE as described above. Gels were scanned at 700 nm (red channel for Cy5 detection) as described previously and data were processed in Image Studio software. Each fluorescence band was quantified, and enzyme inhibition profiles were created in GraphPad Prism 7 software. Fluorescent band with no TOF-probe (0 μM) served as control (0% inhibition), and dark black/background band showed no protease activity (100% inhibition). In a similar manner, TOF-probes’ selectivity was assessed in THP-1 cells. In brief, 100,000 of non-adherent THP-1 cells were placed into 12-well plates, and incubated with various concentrations of TOF-probes, followed by additional incubation with Cy5-labeled probes. For THP-1 only cathepsin L and cathepsin B Cy5 ABPs were used, as this cell line display almost no legumain activity. All experiments were performed in duplicate and data on graphs was presented as averages.

### Flow cytometry analysis

50,000 of HCT-116 and MDA-MB-231 cancer cells were seeded in 12-well plates and allowed to attach overnight. In parallel, 50,000 of non-adherent THP-1 cells were placed in 12-well plates. The following day, 1 μM Cy5-labeleld ABPs for cathepsin L (MP-cL3), cathepsin B (MP-cB2) and legumain (Cy5-MP-L01) were added and incubated with cells for four hours. Control samples were pre-incubated with E64D (25 μM, 2 hours). Two experiments were performed: (1) all three probes were incubated with cells and (2) in each well one probe was added and after four hours of incubation, samples were combined prior to flow cytometry analysis. After incubation, cells were washed twice with DPBS and harvested. In case of separate ABPs incubation, cells from three wells (cathepsin B, cathepsin L and legumain) were combined. Cells were then pelleted by centrifuging at 500 x *g*, washed with DPBS and fixed with 4% PFA in DPBS for 20 minutes at 4°C. Next, samples were washed once with DPBS, re-suspended in CyFACS buffer and kept on ice until flow cytometry acquisition, but no longer than one hour. Multicolor flow cytometry was performed with single cell suspension (10^6^ cells) on LSRFortessa (14 color) at Sanford Burnham Prebys Flow Cytometry Core Facility and 10,000 events were collected for each sample. Data were analyzed using FlowJo software.

### Mass cytometry analysis of cancer cell lines with the use of cysteine protease TOF-probes

In order to apply our TOF-probes in Helios mass cytometry system, 200,000 cells/well of each cell line (HCT-116 and MDA-MB-231) were seeded onto 12-well plate and allowed to attach overnight. In parallel, 50,000 of non-adherent THP-1 cells were placed in 12-well plates. The next day, medium was exchanged and 2 μM TOF-probes were incubated with cells for four hours. After this time, cells were harvested for mass cytometry analysis. As sample preparation prior to CyTOF acquisition requires multiple washes, cell loss is greater than in case of flow cytometry. Therefore, the cells incubated with three probes were prepared in duplicate and combined after incubation in order to obtain more cells. In case of separate incubation three wells were combined, each containing cells stained with different TOF-probes. Cells were placed in 1.5 mL Eppendorf tubes, pelleted via 5 min centrifugation at 500 x g and washed once with DPBS. Next, cells were fixed with the use of 4% PFA/DPBS for 20 minutes at 4°C and washed again with DPBS. In order to stain DNA, samples were pelleted, supernatant was aspirated and cells were first washed once with 1xPermeabilization Buffer and then re-suspended in 100 μL of 250 nM Ir191/193 intercalator/1xPermBuffer. After 30 minutes of incubation at room temperature samples were spun down, supernatant was aspirated, cells were re-suspended in 500 μL of CyFACS and kept in the fridge until CyTOF acquisition, but no longer than 12 hours. Prior to acquisition, cells were washed twice with CyFACS buffer and twice with diH_2_O.

### Peripheral blood mononuclear cells analysis

Fresh blood was isolated from a healthy donor (60 mL) and transferred to storage vessel. An equal volume of 2% FBS/DPBS was added to blood and mixed by swirling. Next, four 50 mL conical tubes were filled with 15 mL of Lymphoprep™ and 30 mL of blood was carefully layered on top of Lymphoprep™. Tubes were centrifuged at 800 x *g* for 30 minutes with the deceleration set to 1. After centrifugation the PBMCs was transferred to a sterile 50 mL falcon tube, an equal volume of 2% FBS/PBS was added and mixed carefully. Tubes were centrifuged at 400 x *g* for 5 minutes and the supernatants were aspirated. Cells were re-suspended in 45 mL of RBC Lysis Buffer (diluted at 4.5 mL of lysis buffer in 40.5 mL H2O) and incubated at room temperature for 7 minutes followed by centrifugation (400 x *g* for 5 minutes). Supernatant was aspirated, and cells were washed with 2% FBS/PBS, centrifuged using the same settings, and re-suspended in 45-50 mL RPMI + 10% FBS + antibiotics (100 units/mL penicillin and 100 μg/streptomycin). Cells were counted and 50,000 cells per sample were pelleted, re-suspended in media containing 2 μM TOF-probes (various sets of probes, incubated together or separately) and plated onto 12-well dishes. Fresh PBMC’s were incubated with TOF-probes for six hours, harvested and washed twice with DPBS. Next, each sample was fixed with 4% PFA/DPBS for 20 minutes at 4°C and washed once with DPBS and once with CyFACS buffer. Primary antibodies were purchased from Fluidigm^®^ and used as described below. Antibody cocktail was prepared by adding 1 μL of each pre-titrated antibody (anti-hCD3, CD19, CD56, CD45, CD11c, and CD14) into 50 μL of CyFACS buffer. Next, cells were re-suspended in 50 μL of CyFACS buffer and 50 μL of antibody cocktail was added to each sample. Cells were mixed carefully by pipetting and incubated in the cocktail for one hour on ice. After this time, cells were pelleted (800 x *g*), supernatant was aspirated, cells were washed once, centrifuged and re-suspended in 500 μL of CyFACS buffer. Samples were kept in the fridge (4°C) overnight. The following day, cells were pelleted and re-suspended in 100 μL of 1x Permeabilization Buffer (ThermoScientific) containing 250 nM Ir191/193 DNA intercalator. After 30 minutes of incubation, samples were washed twice with CyFACS buffer, twice with ddH_2_O and kept on ice until CyTOF acquisition (no longer than 2 hours). Prior to CyTOF run, pellet was adjusted to 2.5-5 × 10^5^/mL. Data was acquired on a CyTOF (Fluidigm^®^). Obtained data were converted to an FCS file and analyzed by FlowJo software (Treestar Inc.).

### Detection of active proteases using Hyperion Imaging System

To visualize active cathepsin B, cathepsin L and legumain in THP-1 cells we employed Hyperion imaging mass cytometry system. Customized three-well adhesion slides were purchased from Marienfield, Germany. THP-1 cells were suspended in DBPS, attached on slides (20,000 cells/well) and left for an hour in the incubator at 37°C until adhered. Next, cells were washed once with DPBS and fixed with 100 μL of 2%PFA/DPBS for 20 minutes at room temperature. Slides were washed twice with DPBS in coplin jar and cells were permeabilized with 100 μL of 0.1% Triton-X for 10 minutes. Again, slides were washed twice with DPBS followed by blocking with 100 μL of 1% BSA in PBS and incubated for 40 minutes at room temperature in order to avoid unspecific binding of the probes. Afterward, THP-1 cells were washed twice and 100 μL of 2 μM probe preparation was incubated for four hours in 37°C, 5% CO_2_. Cells were then washed with DBPS and 250 nM Ir191/193 solution was added in order to label DNA within the cells. After 30 minutes of incubation, cells were again washed, rinsed with water for 5 seconds and air dried at room temperature. The prepared slides were subjected to IMC analysis, respectively. At least three Region of Interests (ROIs) of area 400*400 mm were ablated with a laser frequency 200 Hz and ion count was measured by CyTOF Helios mass cytometer as described earlier [22]. DNA intercalator ^191^Ir/^193^Ir was used for cell segmentation.

## Supporting information

Supplemental Section

## ACKNOWLEDGMENTS

This project has received funding from the European Union’s Horizon 2020 research and innovation program under the Marie Skłodowska-Curie grant agreement No. 661187 (to MP), and from National Institutes of Health in the USA (grant GM99040 to GSS), and from Sanford Burnham Prebys NCI Cancer Center Support Grant P30CA030199 (to GSS). Marcin Drag lab is supported by the Foundation for Polish Science, and National Science Centre in Poland (grant Harmony UMO-2018/30/M/ST5/00440). Boris Turk research is supported by Slovene Research Agency (gran P1-0140). We thank Yoav Altman, a director of the Flow Cytometry Facility at SBP Medical Discovery Institute, for his support with flow cytometry analysis. WR is a beneficiary of a START scholarship from the Foundation for Polish Science.

