## Supplemental Section for "The Activome: multiplexed probing of activity of proteolytic enzymes using mass cytometry-compatible activity-based probes (TOF-probes)"

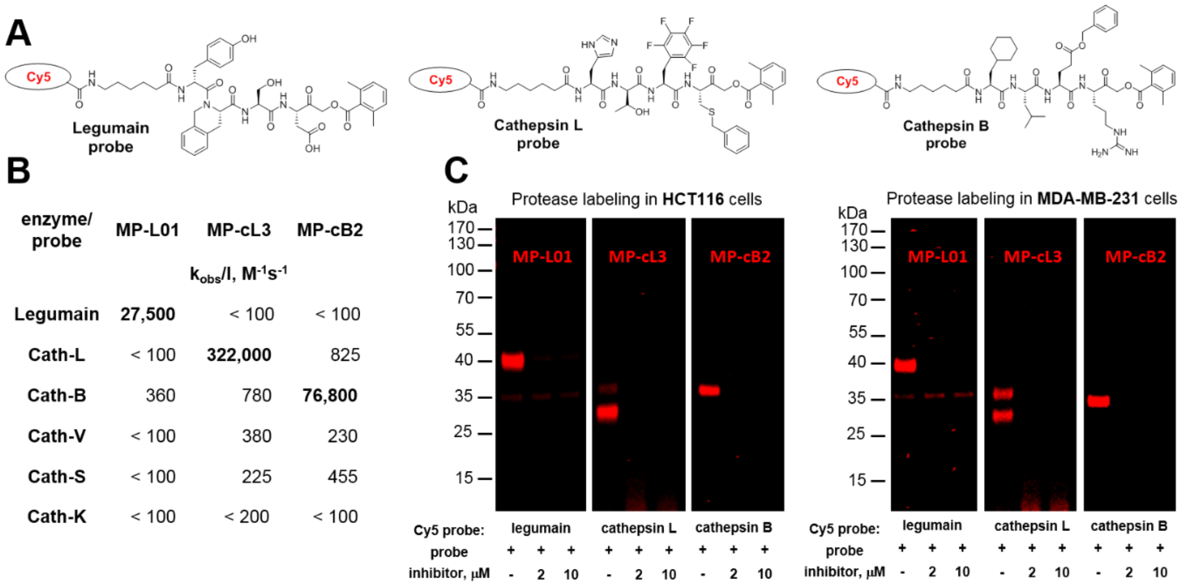


**Supplemental Figure 1** Selective Cy5-labeled ABPs for the detection of lysosomal proteases in cancer cells. **A** Structures of three selective Cy5-labeled ABPs for legumain, cathepsin L and cathepsin B. **B** Kinetic parameters (second order inhibition constant, $k_{obs}/[I]$) of ABPs. **C** Selective labeling of active legumain, cathepsin L and cathepsin B in cancer cells lines (HCT-116 and MDA-MB-231) with Cy5-ABPs. Probes (1 µM) were incubated with cells for 4 hours. For enzyme inhibition MP-L01 (for legumain) and E64d (for cathepsins) were incubated for 2 hours prior to enzyme labeling with ABPs.

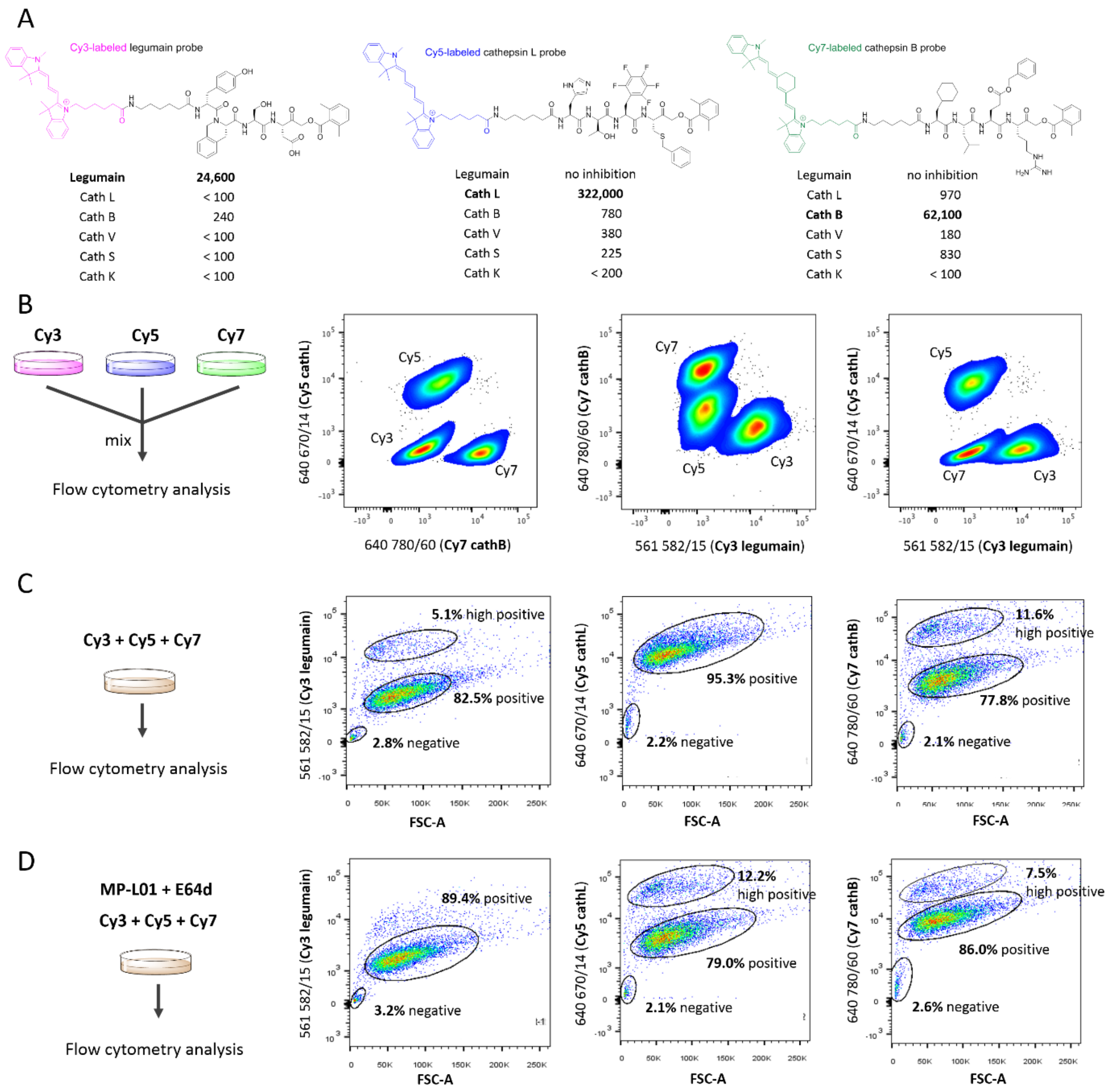


**Supplemental Figure 2** Detection of legumain and cathepsins in HCT116 cancer cells with fluorescent probes and flow cytometry. **A** Structures and inhibition constants ($k_{obs}$/[I], $M^{-1}s^{-1}$) of three selective ABPs for legumain (Cy3-labeled), cathepsin L (Cy5-labeled) and cathepsin B (Cy7-labeeld). **B** HCT-116 cells were incubated with three ABPs separately, then combined and subjected for flow cytometry. **C** HCT-116 cells were incubated with a mixture of three ABPs followed by flow cytometry analysis. Data from both experiments are presented on separate two-dimensional plots. **D** HCT-116 cells were pre-incubated with legumain (MP-L01) and pan-cathepsin (E64d) inhibitors followed by incubation with a mixture of three ABPs. Data are presented on separate plots where the x-axis show forward scatter area (FSC-A), and y-axis shows the intensity of ABP`s fluorescent signal.

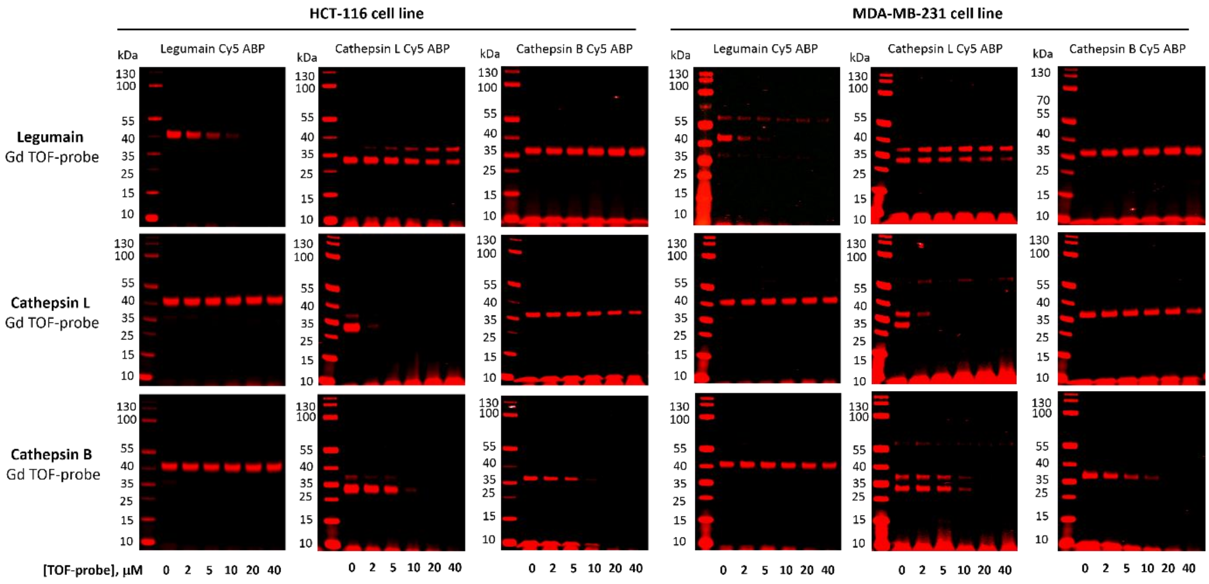


**Supplemental Figure 3** Whole gels demonstrating the potency and selectivity of TOF-probes for the labeling/inhibition of legumain, cathepsin L and cathepsin B in human cancer cell lines, HCT-116 and MDA-MB-231. To assess the selectivity of TOF-probes, they were incubated at various concentration range with cancer cells, and next the residual protease activity was detected with selective, Cy5-labelled probes (Cy5-MP-L01 for legumain, Cy5-MP-cL3 for cathepsin L and Cy5-cB2 for cathepsin B).

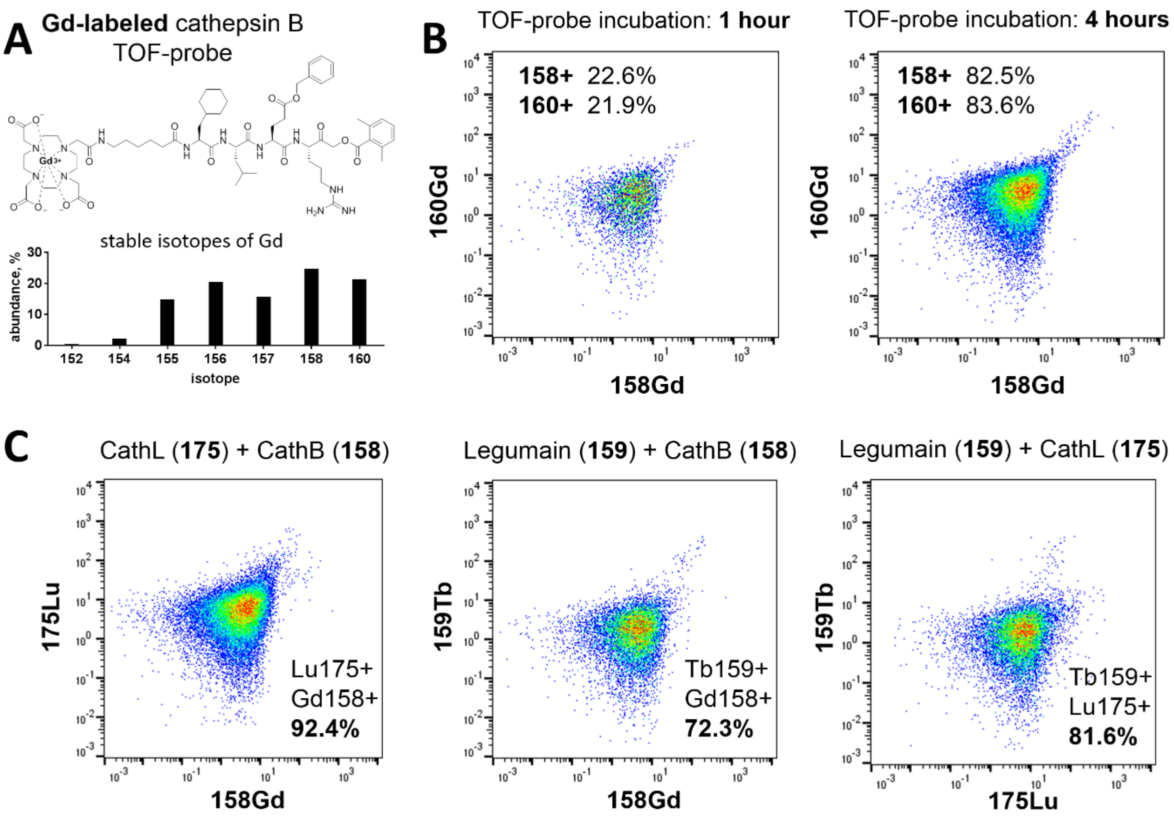


**Supplemental Figure 4** Selective detection of legumain, cathepsin L, and cathepsin B in HCT-116 cell line. **A** The structure of cathepsin B TOF-probe labeled with Gd metal. By chelating MP-cB2 probe with non-isotopically pure Gadolinium metal we detected cathepsin B activity on two channels (158 and 160) and demonstrated the homogenous distribution of TOF-probe within single cells. **B** Cathepsin B Gd TOF-probe uptake kinetic in HCT-116 cells. After 1 hour of incubation only around 20% of cells were $^{158}$Gd and $^{160}$Gd positive, whereas the prolonged incubation (4 hours) resulted in the labeling of cathepsin B in over 80% of cells population. Symmetrical graphs demonstrated that TOF-probe was homogenously distributed. **C** Parallel detection of three proteases in HCT-116 cells. Three TOF-probe were incubated with cells for 4 hours and subjected for mass cytometry analysis. Results demonstrate that all three probes were taken up by cells and efficiently labeled proteases.

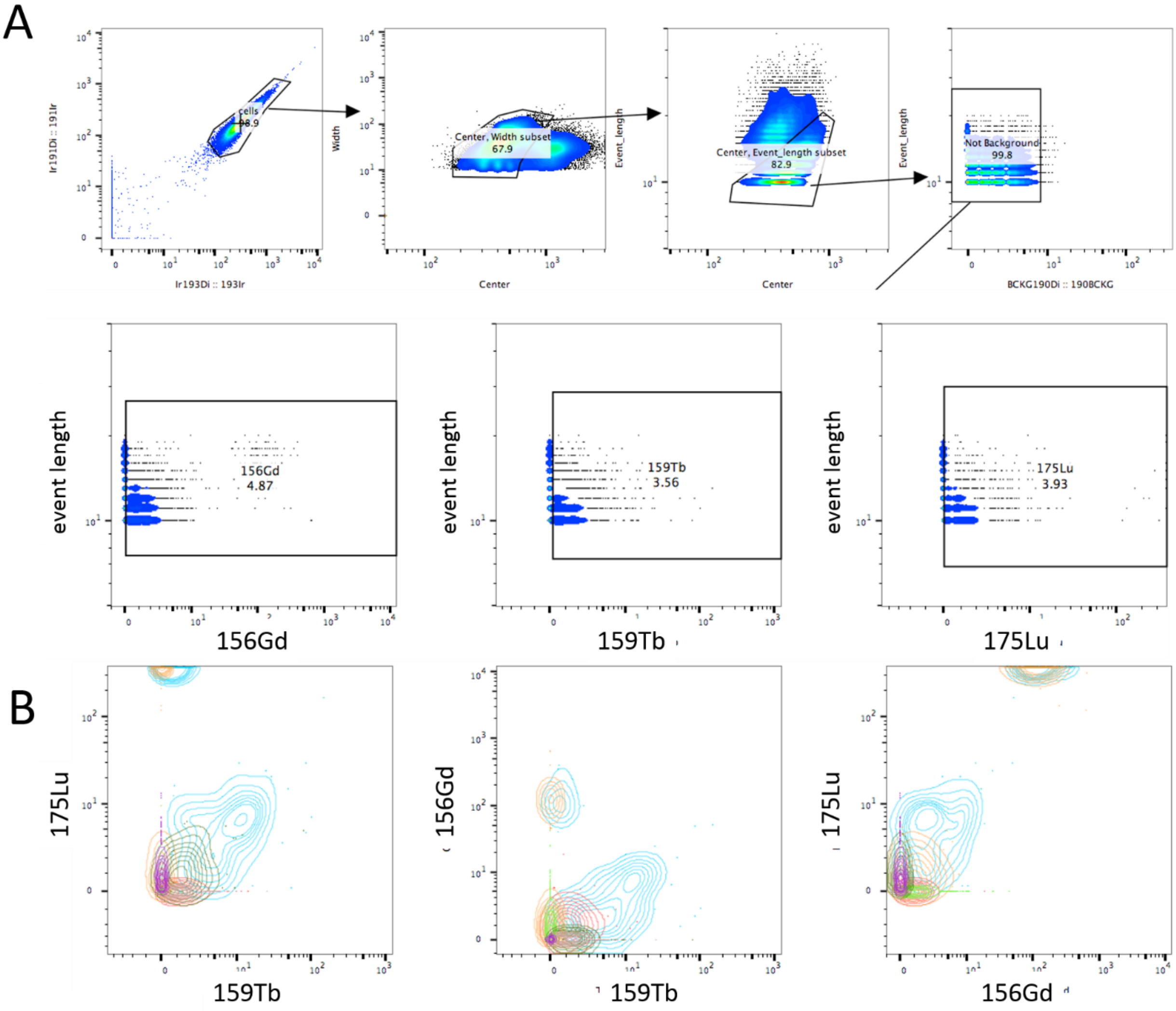


| | Sample Name | Subset Name | Count | Freq. of Parent |
|---|---|---|---|---|
| | HCT116_control_01_01.FCS | 156Gd-159Tb-175Lu- | 49347 | 88.9 |
| | HCT116_control_01_01.FCS | 156Gd-159Tb-175Lu+ | 1758 | 3.17 |
| | HCT116_control_01_01.FCS | 156Gd-159Tb+175Lu- | 1603 | 2.89 |
| | HCT116_control_01_01.FCS | 156Gd-159Tb+175Lu+ | 106 | 0.19 |
| | HCT116_control_01_01.FCS | 156Gd+159Tb-175Lu- | 2268 | 4.09 |
| | HCT116_control_01_01.FCS | 156Gd+159Tb-175Lu+ | 164 | 0.30 |
| | HCT116_control_01_01.FCS | 156Gd+159Tb+175Lu- | 118 | 0.21 |
| | HCT116_control_01_01.FCS | 156Gd+159Tb+175Lu+ | 152 | 0.27 |

**Supplemental Figure 5** Detailed gating strategy for non-TOF-probes treated HCT-116 cells. **A** DNA ($^{191}$Ir/$^{193}$Ir) positive events were gated based on their width and length. The signal background as well as doublets were removed form analysis. For each channel analyzed ($^{156}$Gd, $^{159}$Tb and $^{175}$Lu) graph plots were gated according to signal intensity. **B** Metal signals were paired (Lu/Tb, Gd/Tb, Lu/Gd) and presented on two dimensional graphs. Based on signal counts, positive/negative events were calculated and summarized in table.

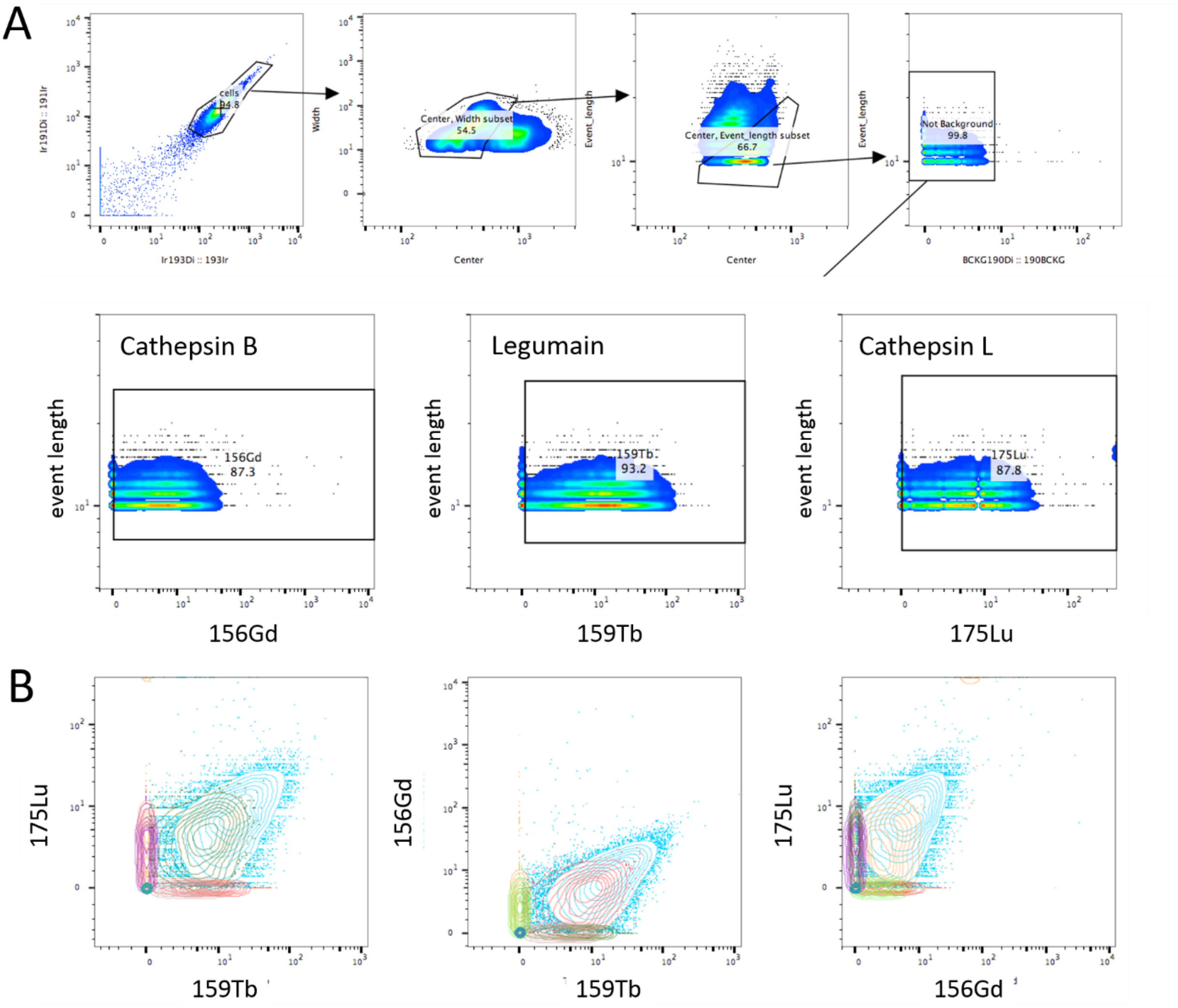


| | Sample Name | Subset Name | Count | Freq. of Parent |
|---|---|---|---|---|
| | HCT_A_2uM_06.FCS | 156Gd-159Tb-175Lu- | 220 | 1.16 |
| | HCT_A_2uM_06.FCS | 156Gd-159Tb-175Lu+ | 255 | 1.34 |
| | HCT_A_2uM_06.FCS | 156Gd-159Tb+175Lu- | 438 | 2.30 |
| | HCT_A_2uM_06.FCS | 156Gd-159Tb+175Lu+ | 1504 | 7.91 |
| | HCT_A_2uM_06.FCS | 156Gd+159Tb-175Lu- | 236 | 1.24 |
| | HCT_A_2uM_06.FCS | 156Gd+159Tb-175Lu+ | 588 | 3.09 |
| | HCT_A_2uM_06.FCS | 156Gd+159Tb+175Lu- | 1422 | 7.48 |
| | HCT_A_2uM_06.FCS | 156Gd+159Tb+175Lu+ | 14356 | 75.5 |

**Supplemental Figure 6** Detailed gating strategy for TOF-probes treated HCT-116 cells exemplified in variant A (Tb probe for legumain, Gd probe for cathepsin B and Lu probe for cathepsin L). **A** DNA ($^{191}$Ir/$^{193}$Ir) positive events were gated based on their width and length, and next the signal background was removed. For each channel analyzed ($^{156}$Gd cathepsin B probe, $^{159}$Tb legumain probe and $^{175}$Lu cathepsin L probe) channels were gated according to signal intensity. **B** Metal signals were paired (Lu/Tb, Gd/Tb, Lu/Gd) and presented on two dimensional graphs. Based on signal counts metal positive/negative events were calculated and summarized in table.

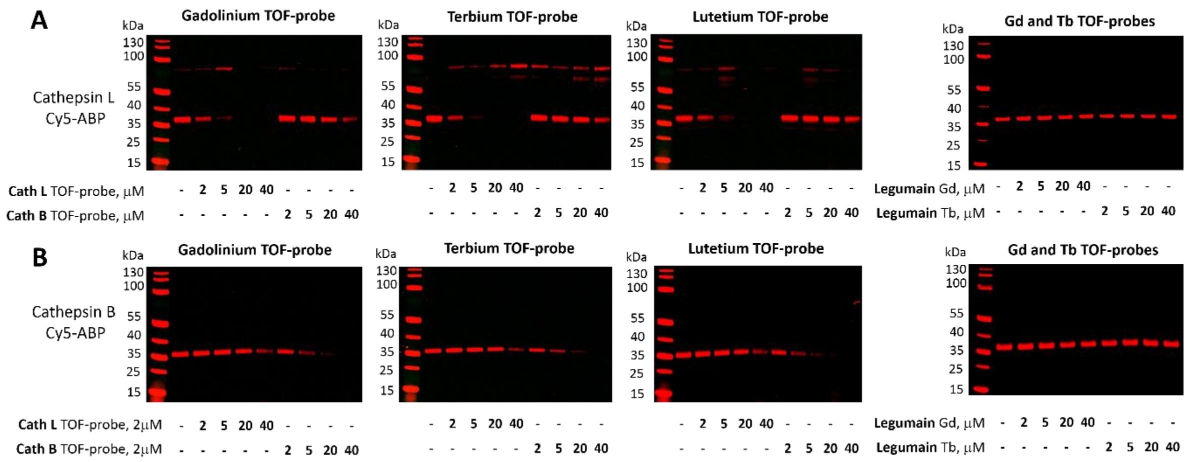


**Supplemental Figure 7** Determination of TOF-probes selectivity toward cathepsin L (A) and cathepsin B (B) in THP-1 cells. To assess the selectivity of TOF-probes, they were incubated at various concentration range with THP-1 cells, and next the residual protease activity was detected with selective, Cy5-labelled probe (Cy5-MP-cL3 for cathepsin L and Cy5-cB2 for cathepsin B).

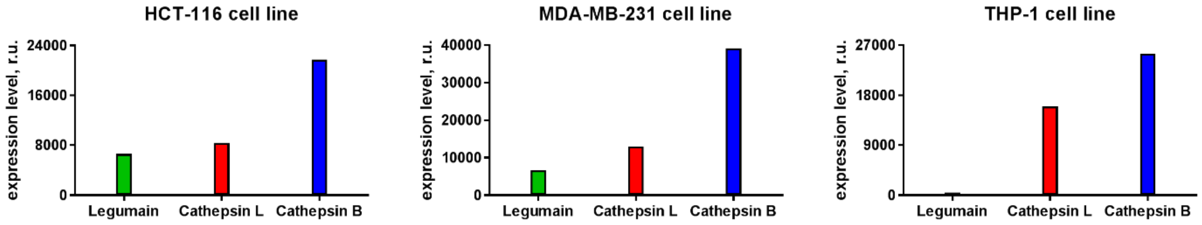


**Supplemental Figure 8** Expression level of legumain, cathepsin L and cathepsin B in three human cell lines: HCT-116, MDA-MB-231 and THP-1. Data was extracted from Geneinvestigator database. Only non-stimulated cells were taken for the analysis, and data presents average expression levels in relative units.

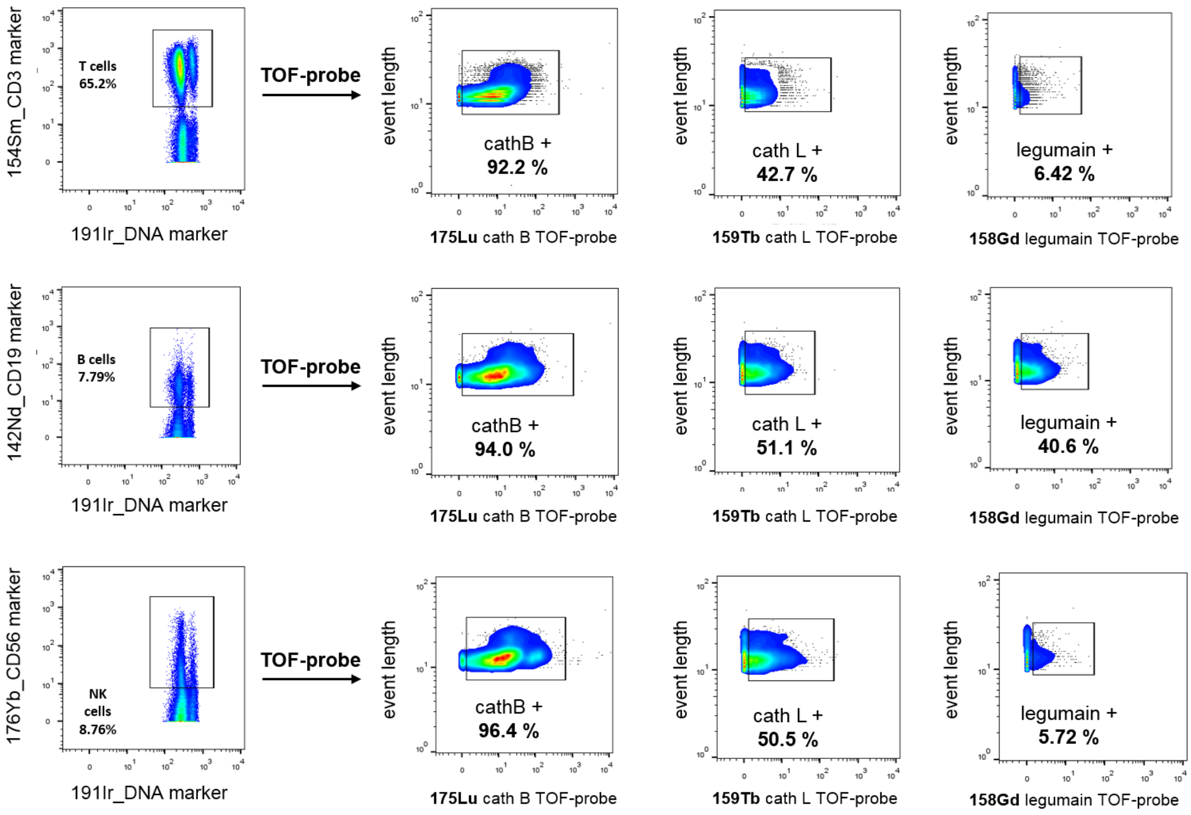


**Supplemental Figure 9** Proteases detection in Peripheral Blood Mononuclear Cells. Different PBMCs` populations were detected with the use of metal-conjugated antibodies ($^{154}$Sm-CD3 for T cells, $^{142}$Nd-CD19 for B cells, and $^{176}$Yb-CD56 for NK cells). Proteases activity was detected with TOF-probes in the following combination: $^{175}$Lu-tagged cat B probe, $^{159}$Tb-tagged cat L probe, and Gd-tagged legumain probe (detected at 158 channel). Data present that cathepsin B is the most active enzyme in all type of cells, cathepsin L is also present in all cell populations, but less active than cathepsin B, and legumain activity is B cells specific.

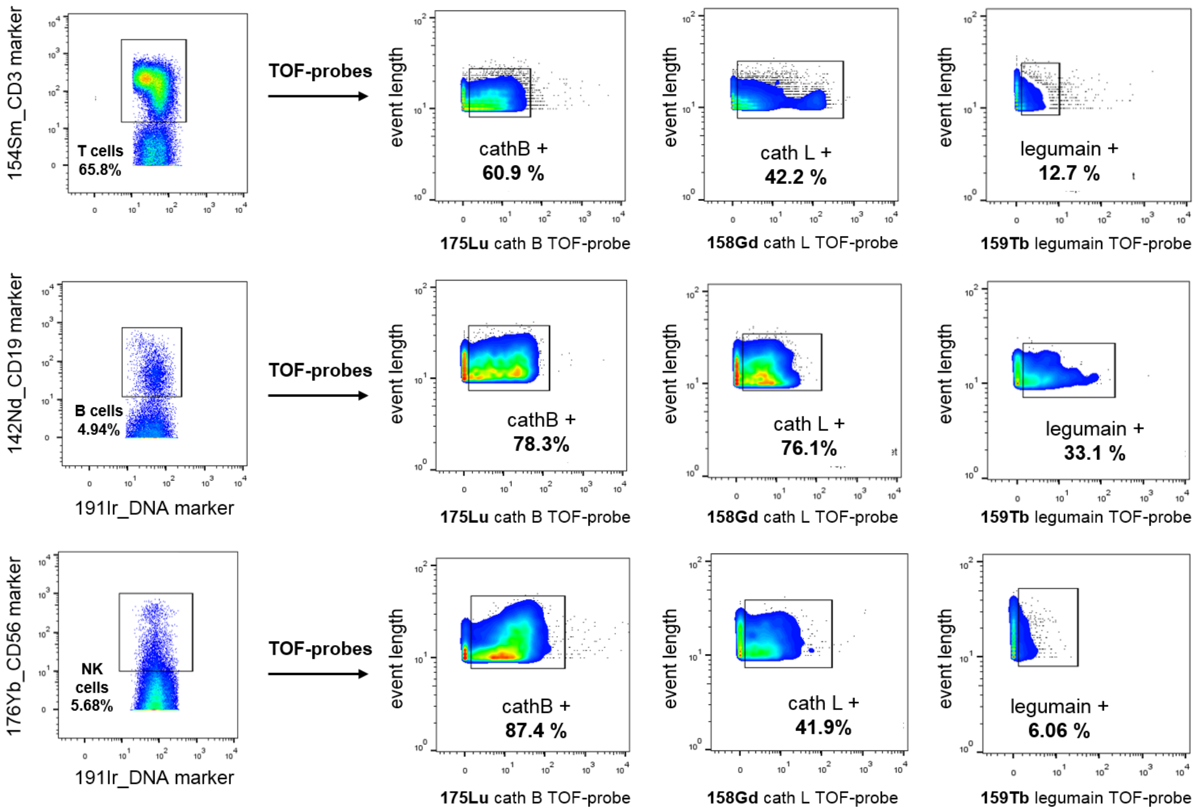


**Supplemental Figure 10** Proteases detection in Peripheral Blood Mononuclear Cells. Different PBMCs` populations were detected with the use of metal-conjugated antibodies ($^{154}$Sm-CD3 for T cells, $^{142}$Nd-CD19 for B cells, and $^{176}$Yb-CD56 for NK cells). Proteases activity was detected with TOF-probes in the following combination: $^{175}$Lu-tagged cat B probe, Gd-tagged cat L probe (detected at 158 channel), and $^{159}$Tb-tagged legumain probe. Data present that cathepsin B is the most active enzyme in all type of cells, cathepsin L is also present in all cell populations, but less active than cathepsin B, and legumain activity is B cells specific.

**DOTA($^{159}$Tb)-Ahx-*D*Tyr-Tic-Ser-Asp-AOMK**

HRMS (m/z): $[M+H]^+$ calcd for $C_{58}H_{74}N_9O_{18}Tb$, 672.7202, found 672.7318

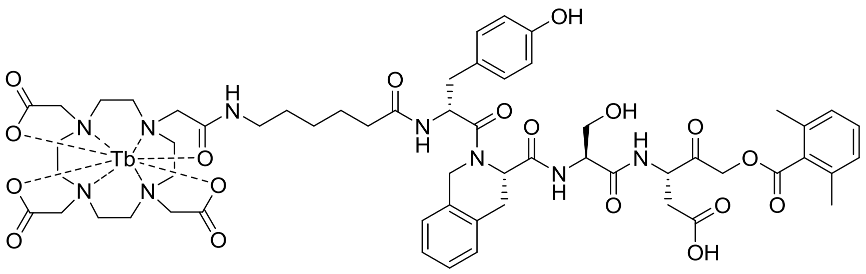

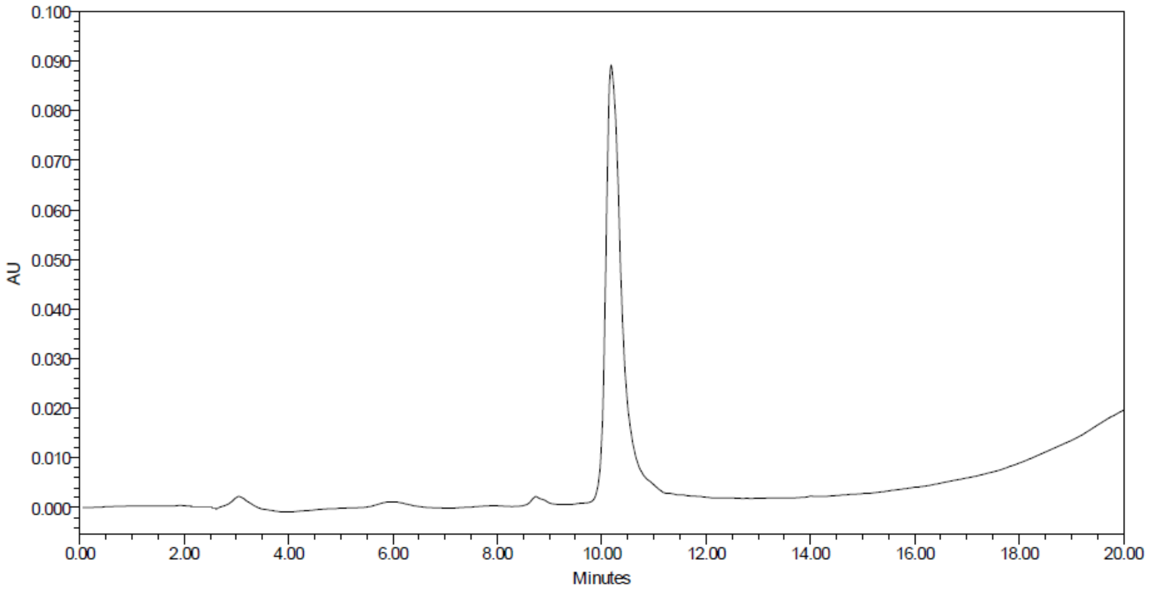


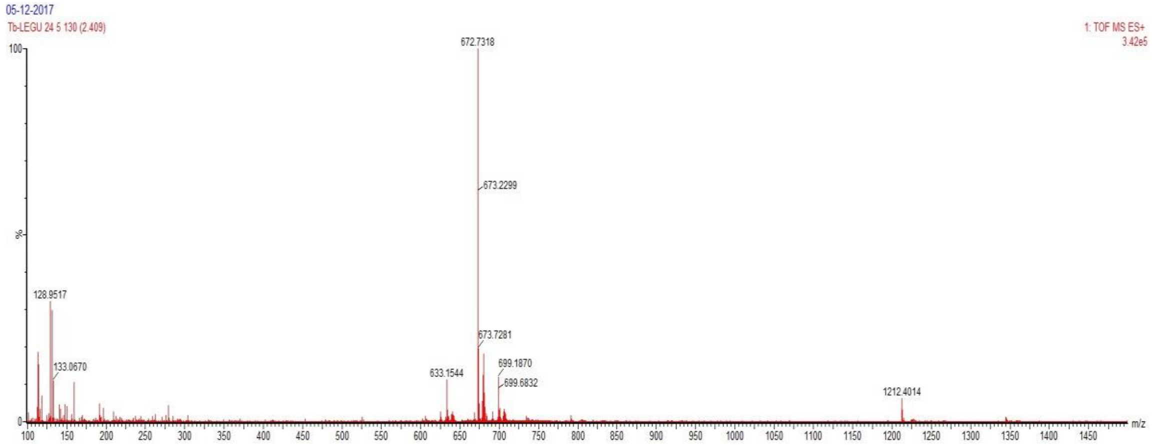

**DOTA(Gd)-Ahx-*D*Tyr-Tic-Ser-Asp-AOMK**

HRMS (m/z): $[M+H]^+$ calcd for $C_{58}H_{74}GdN_9O_{18}$, 1343.4466, found 1343.6244

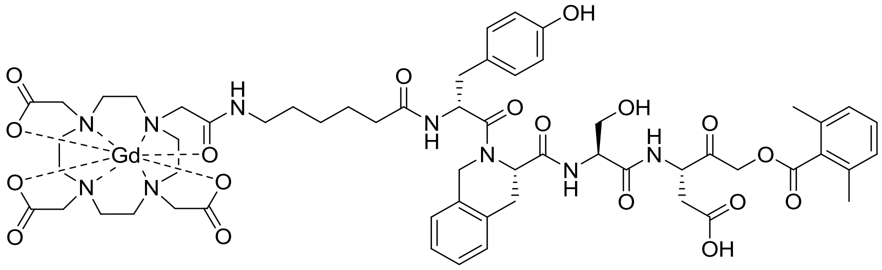

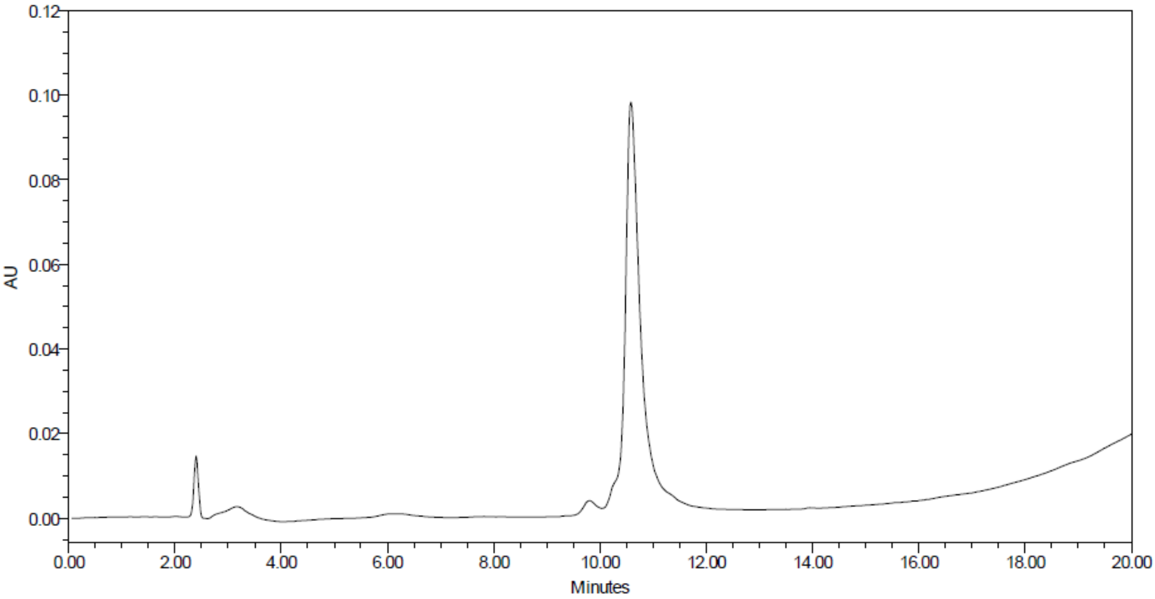


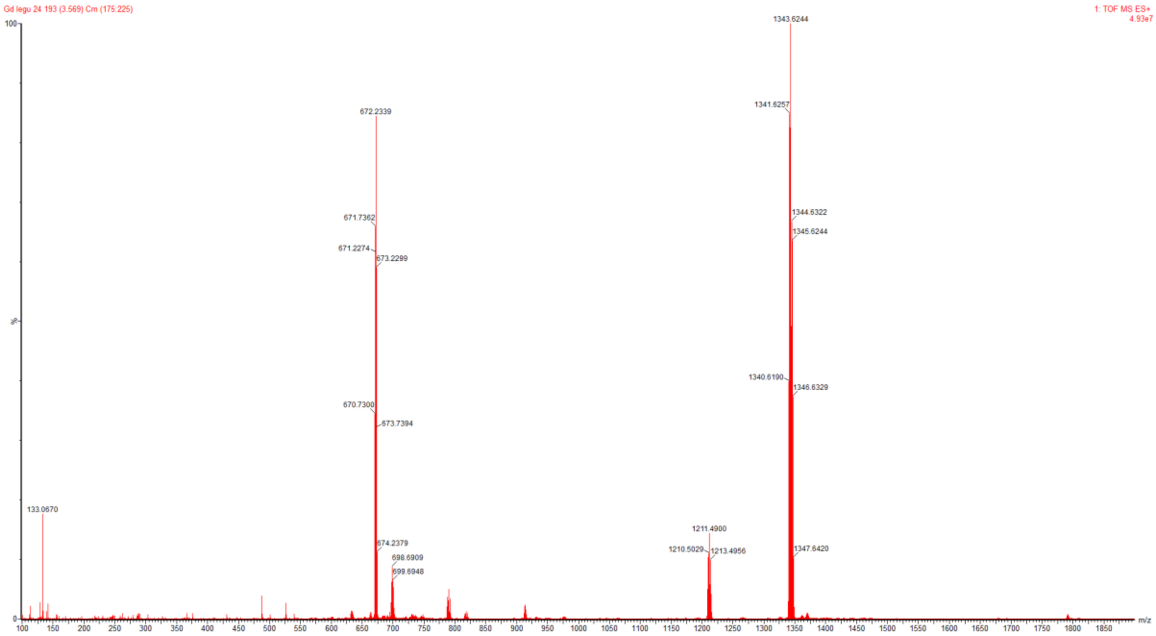

**DOTA([159]Tb)-Ahx-His-*D*Thr-Phe(F5)-Cys(Bzl)-AOMK**

HRMS (m/z): $[M+H]^+$ calcd for $C_{61}H_{75}F_5N_{11}O_{15}TbS$, 1488.4412, found 1488.7352

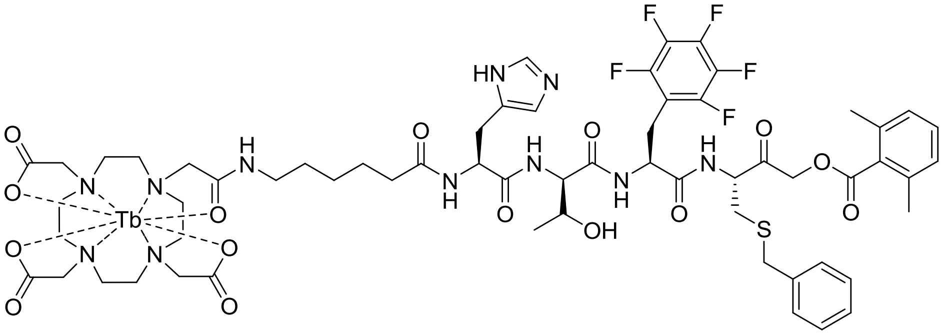

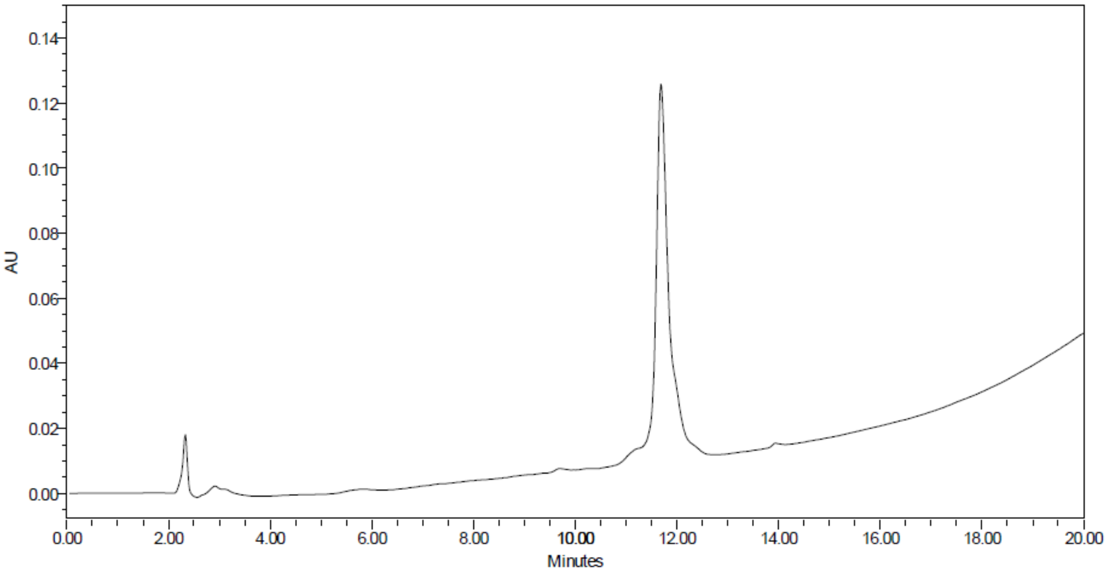


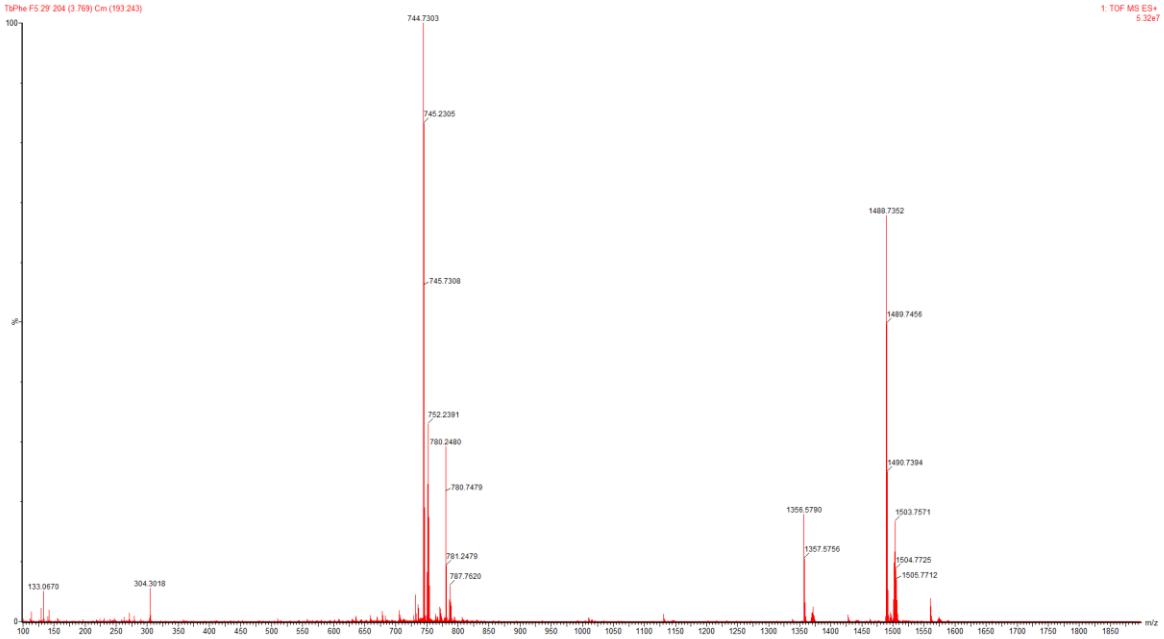

**DOTA($^{175}$Lu)-Ahx-His-*D*Thr-Phe(F5)-Cys(Bzl)-AOMK**

HRMS (m/z): $[M+H]^+$ calcd for $C_{61}H_{75}F_5LuN_{11}O_{15}S$, 1504.4566, found 1504.7725

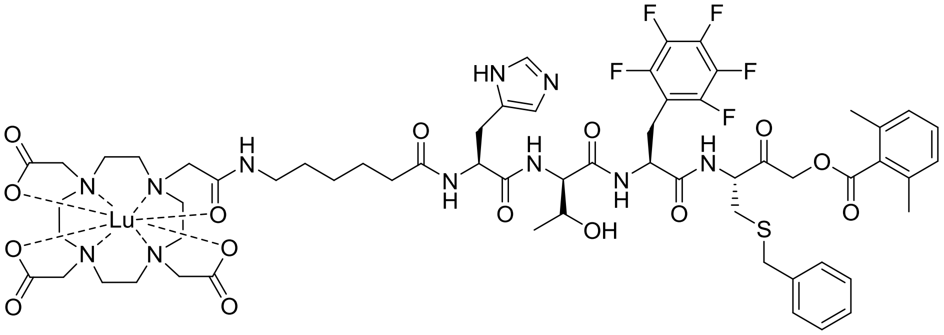

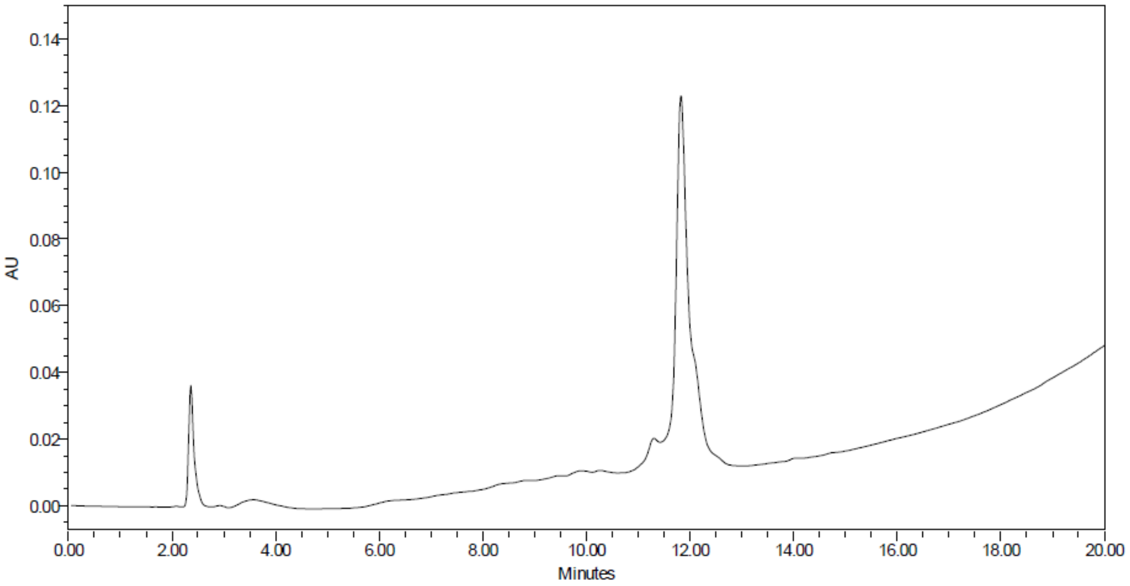

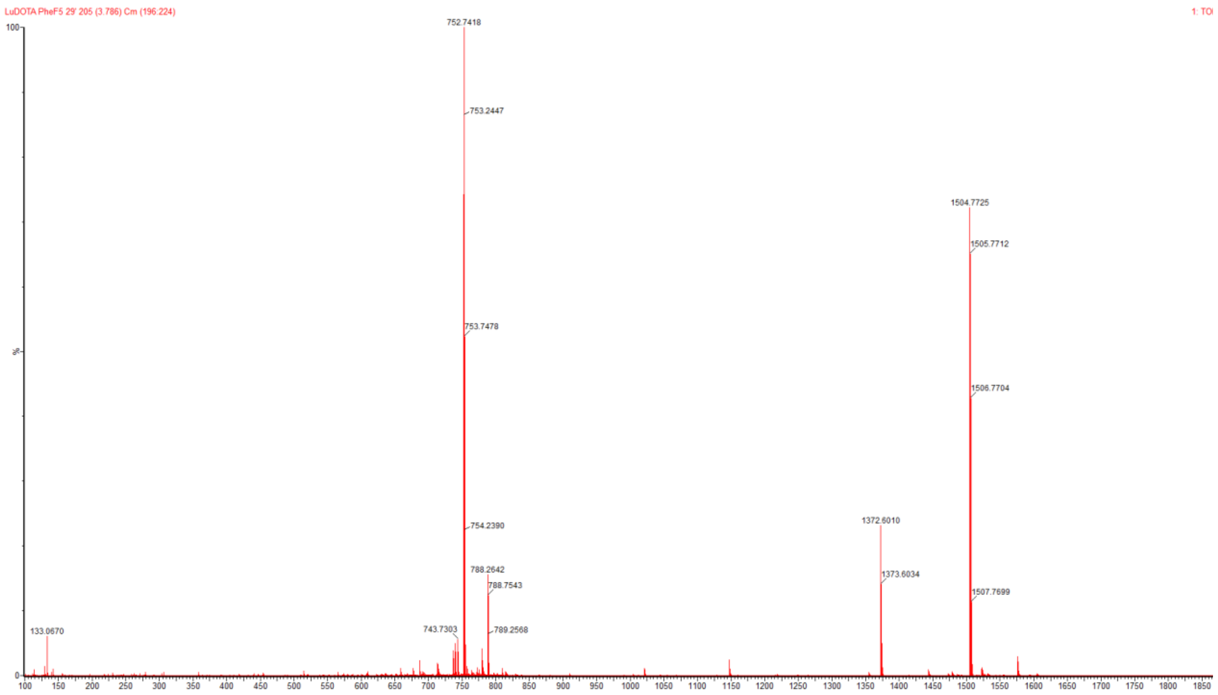


**DOTA(Gd)-Ahx-His-DThr-Phe(F5)-Cys(Bzl)-AOMK**

HRMS (m/z): $[M+H]^+$ calcd for $C_{61}H_{75}F_5GdN_{11}O_{15}S$, 1487.4404, found 1487.4415

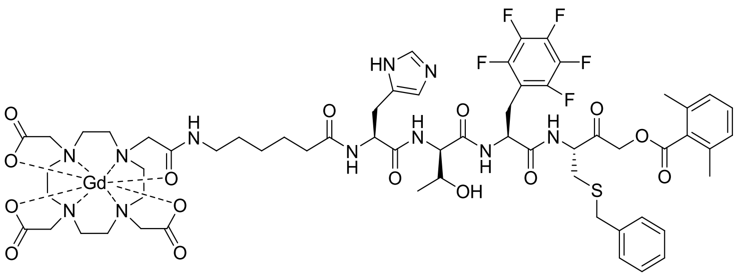

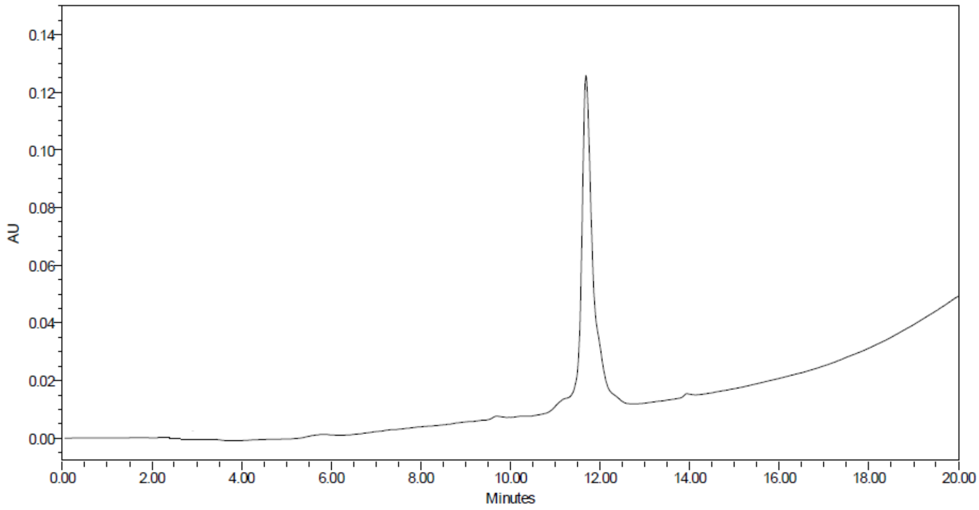

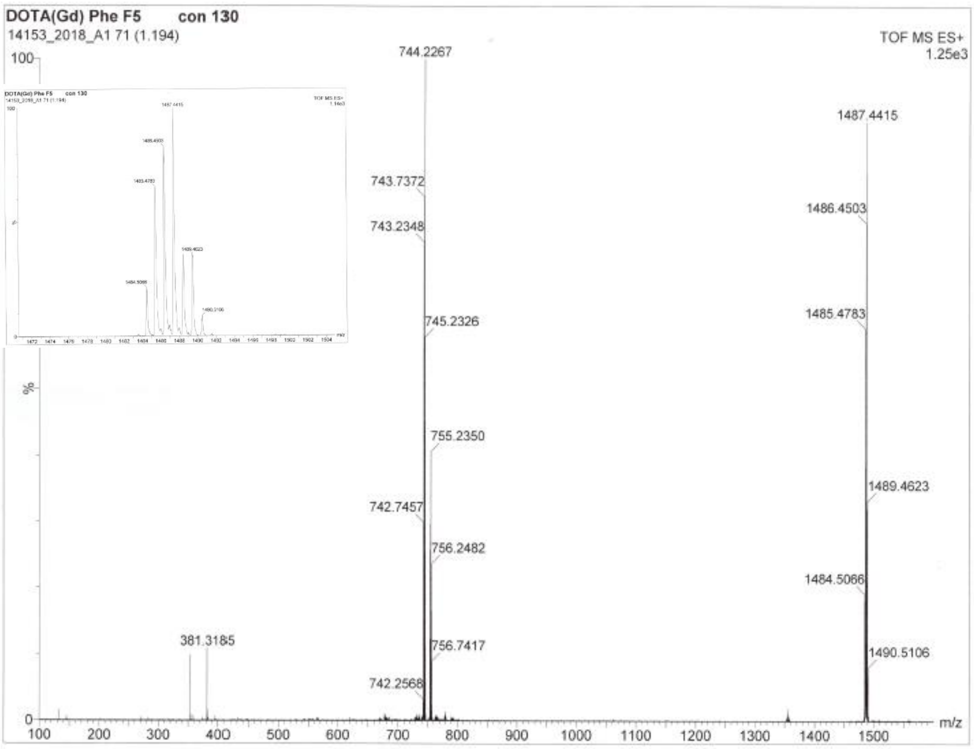


**DOTA($^{159}$Tb)-Ahx-Phe-Cys(Bzl)-AOMK**

HRMS (m/z): $[M+H]^+$ calcd for, $C_{51}H_{66}N_7O_{12}STb$, 1160.3817, found 1160.4764

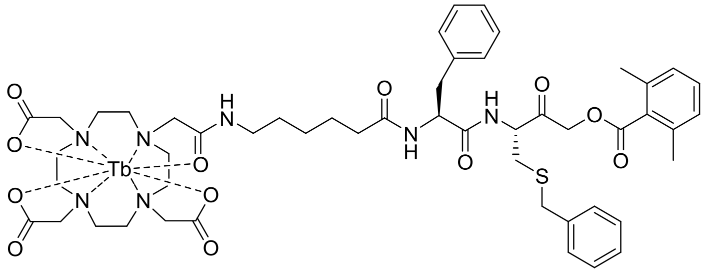

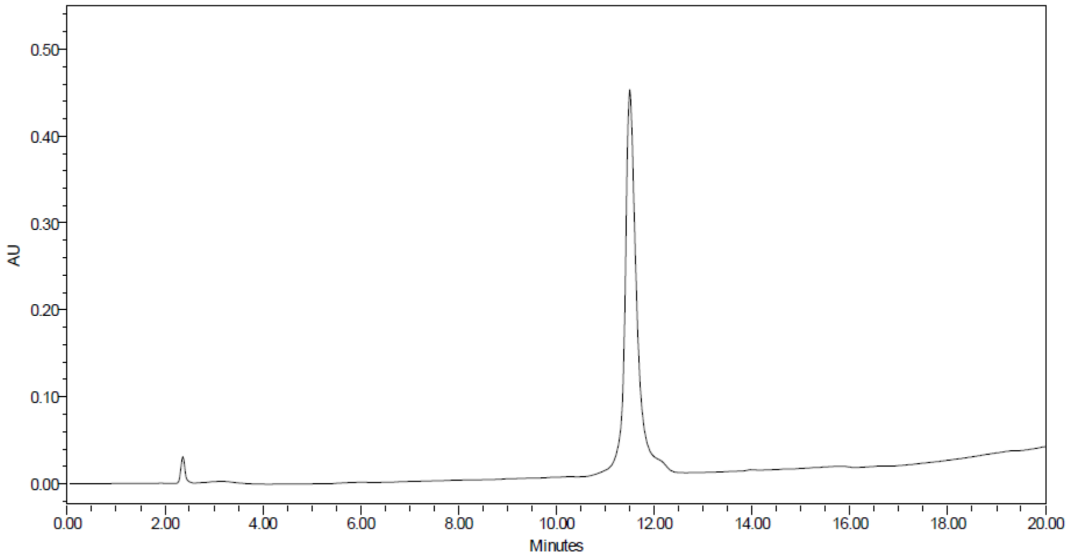


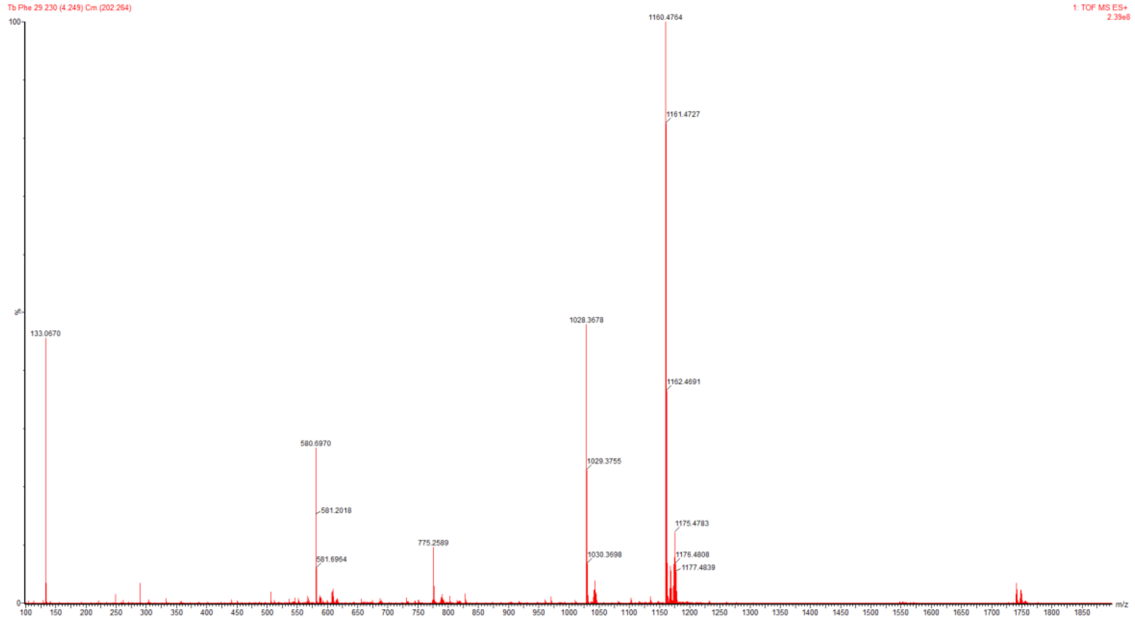


**DOTA($^{175}$Lu)-Ahx-Phe-Cys(Bzl)-AOMK**

HRMS (m/z): $[M+H]^+$ calcd for $C_{51}H_{66}LuN_7O_{12}S$, 1176.3971, found 1176.4958

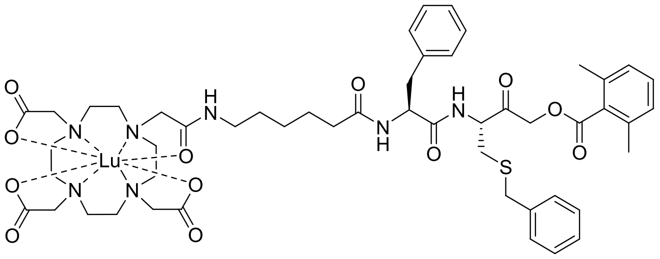

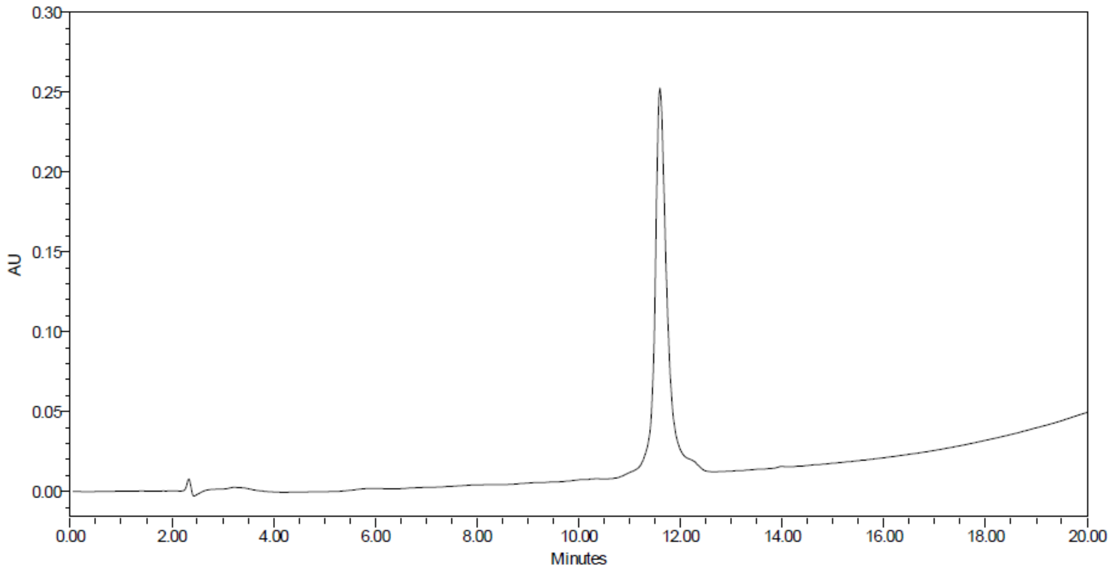
AU
Minutes

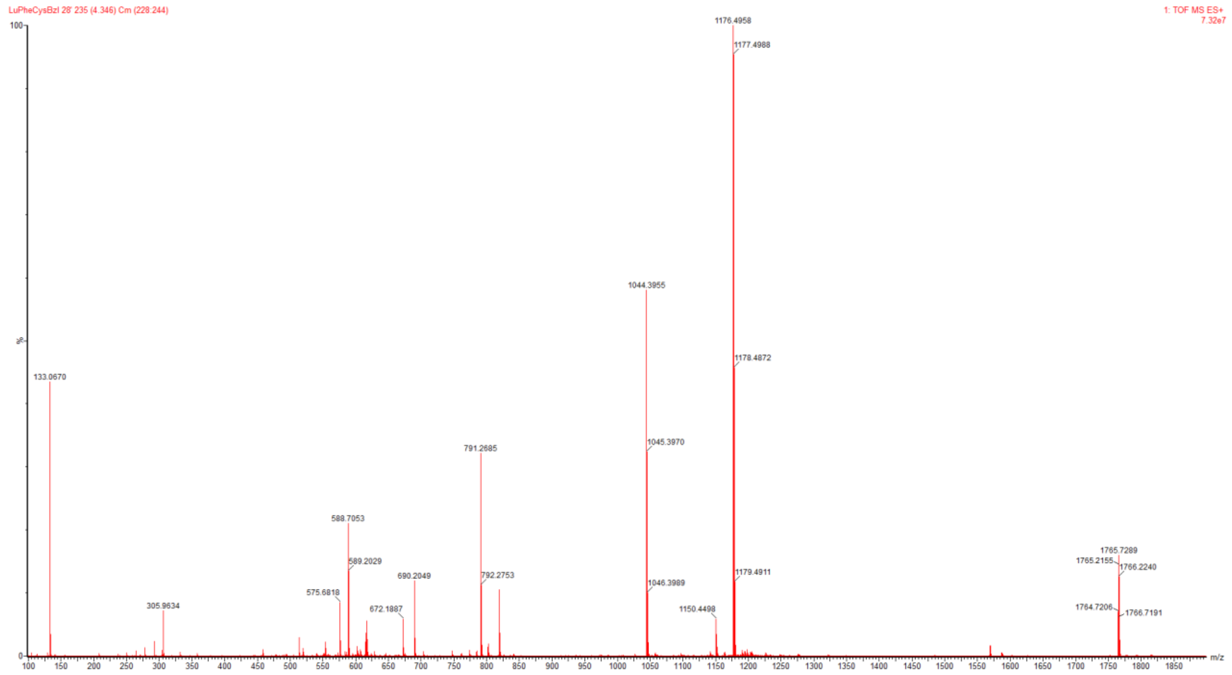
LuPheCysBzl 28' 235 (4.346) Cm (228:244)
1: TOF MS ES+
7.32e7
1176.4958
1177.4988
1178.4872
1179.4911
1044.3955
1045.3970
1046.3989
1150.4498
791.2685
792.2753
588.7053
589.2029
575.6818
690.2049
672.1887
305.9634
133.0670
1765.7289
1765.2155
1766.2240
1764.7206
1766.7191
m/z

**DOTA(Gd)-Ahx-Phe-Cys(Bzl)-AOMK**

HRMS (m/z): $[M+H]^+$ calcd for $C_{51}H_{66}GdN_7O_{12}S$, 1159.3804, found 1159.4659

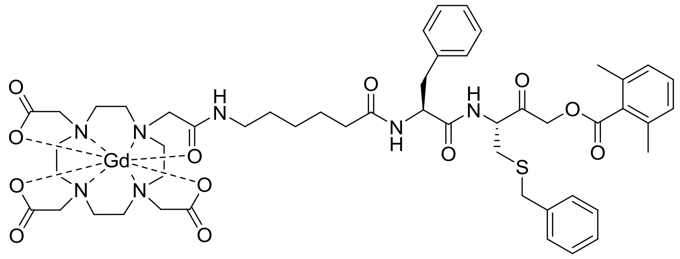

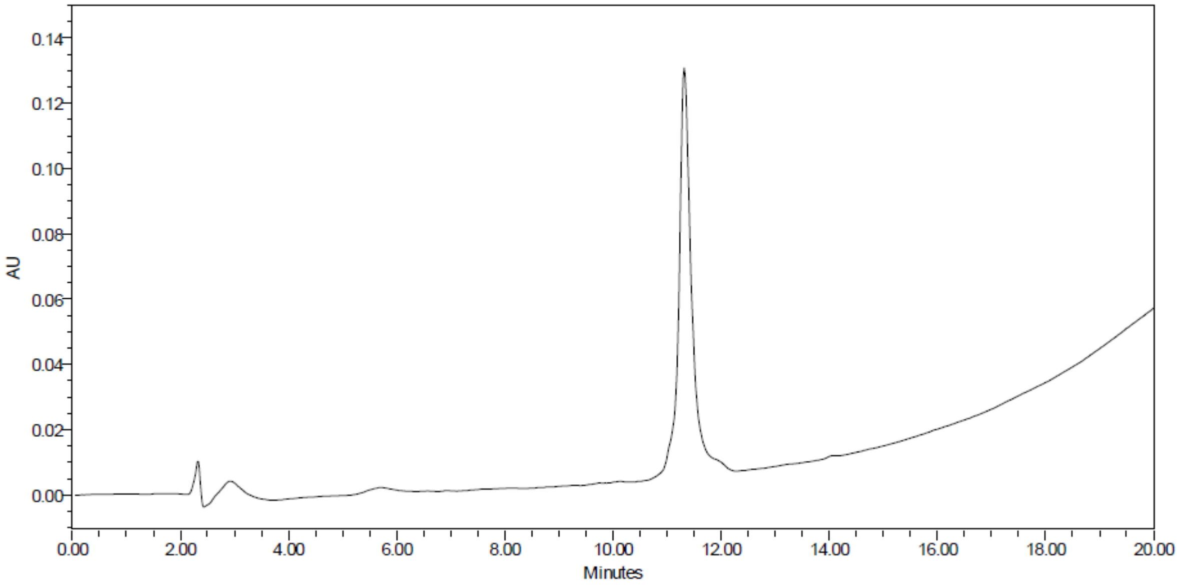


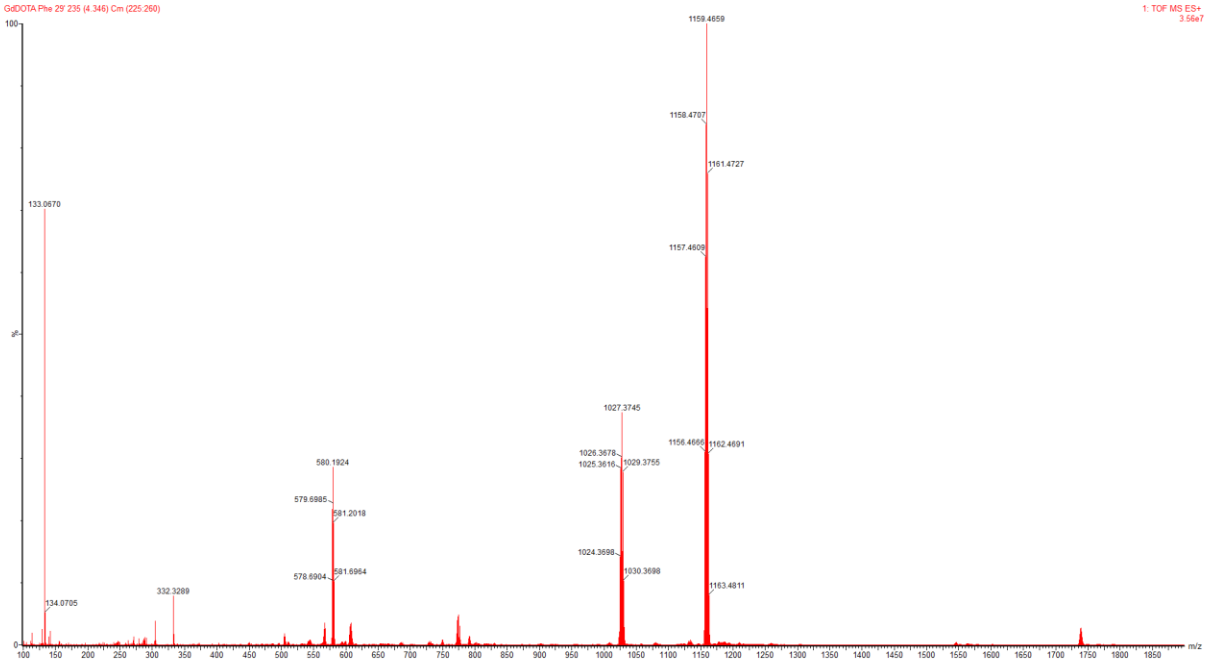

**DOTA(Gd)-Ahx-Cha-Leu-Glu(Bzl)-Arg-AOMK**

HRMS (m/z): $[M+H]^+$ calcd for $C_{65}H_{97}GdN_{12}O_{16}$, 1460.6460, found 1460.9186

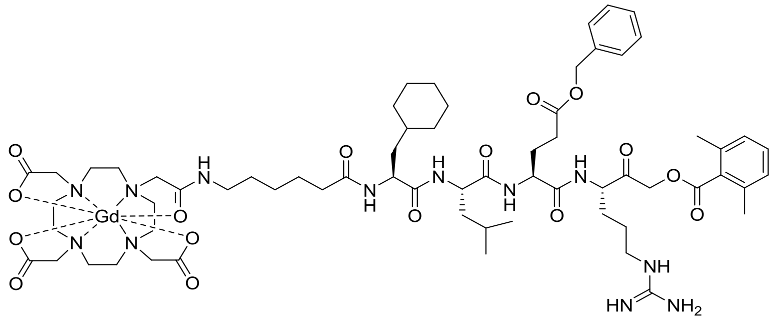

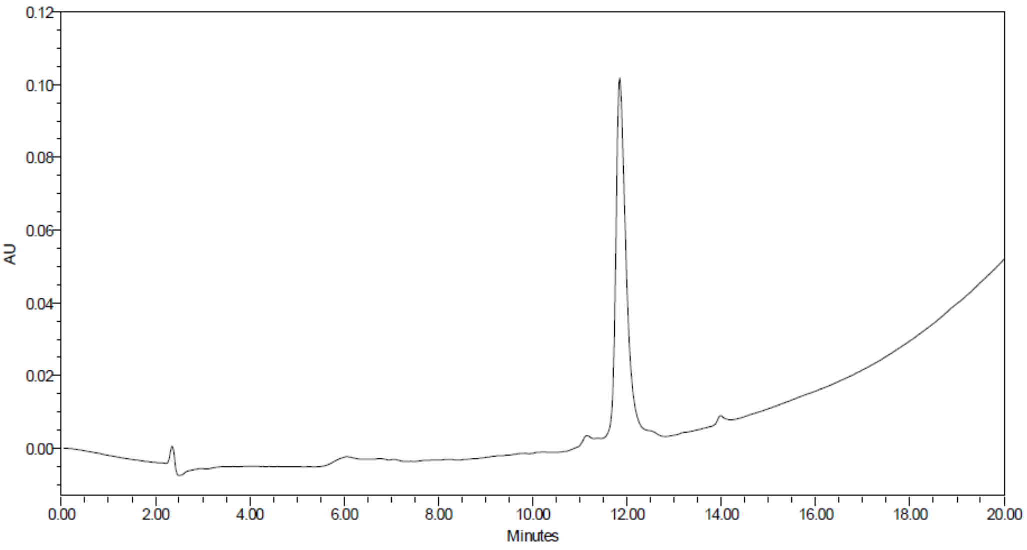


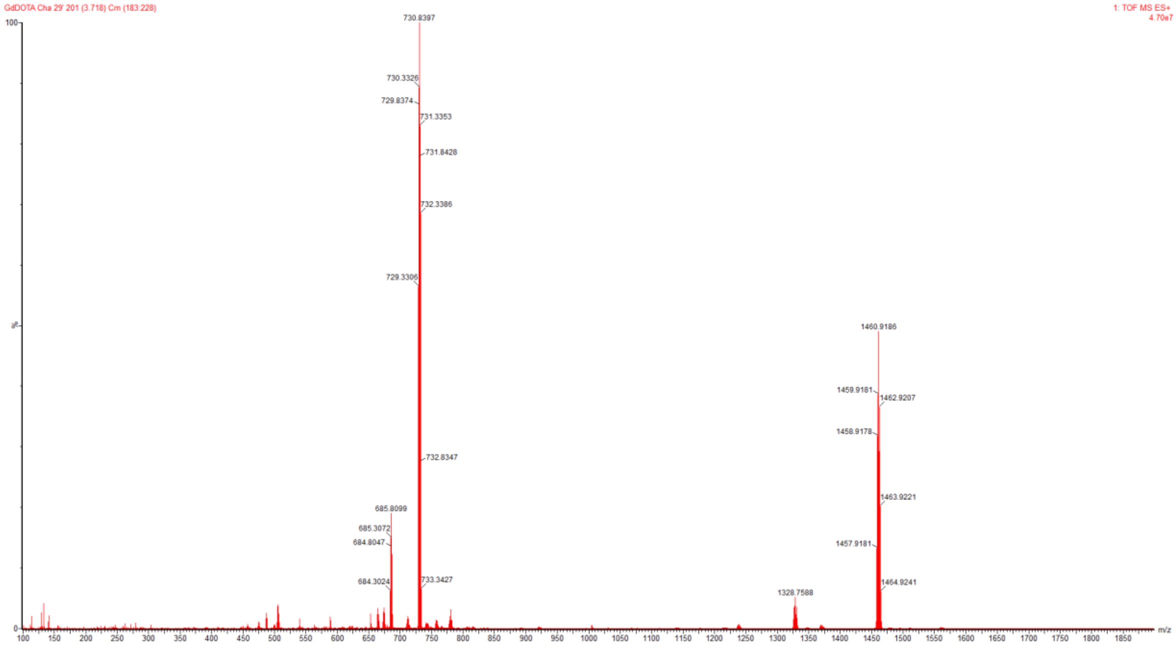

**DOTA($^{159}$Tb)-Ahx-Cha-Leu-Glu(Bzl)-Arg-AOMK**

HRMS (m/z): $[M/2+H]^+$ calcd for $C_{65}H_{97}N_{12}O_{16}Tb$, 731.3273, found 731.3275

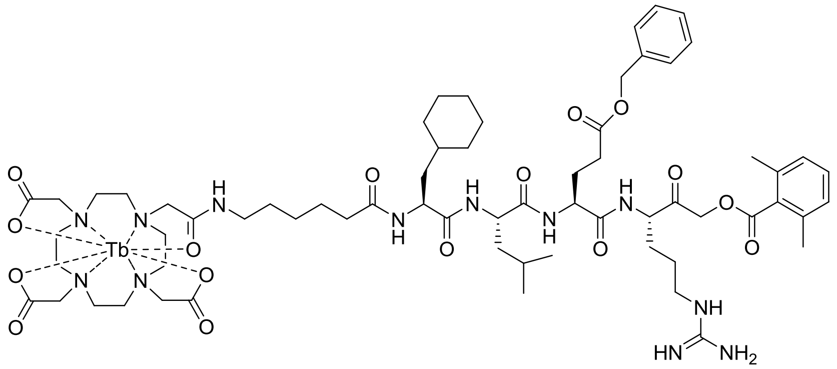


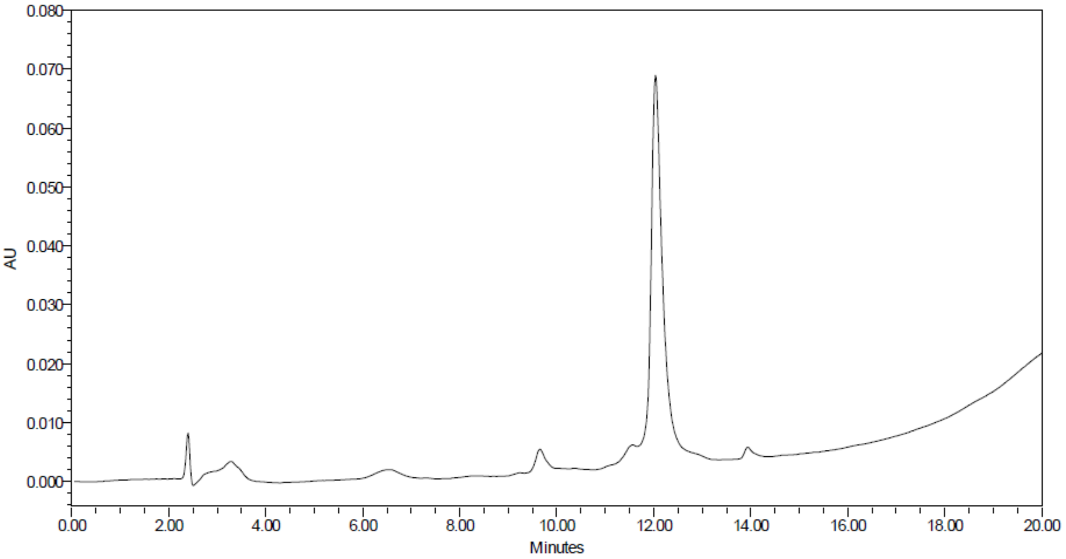

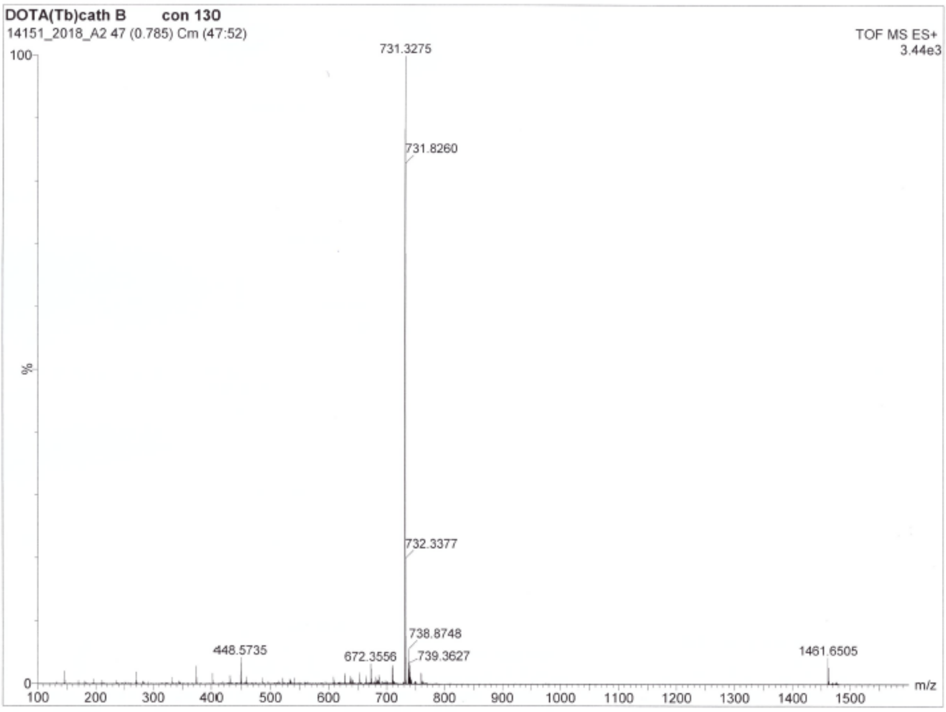


**DOTA($^{175}$Lu)-Ahx-Cha-Leu-Glu(Bzl)-Arg-AOMK**

HRMS (m/z): $[M/2+H]^+$ calcd for $C_{65}H_{97}LuN_{12}O_{16}$, 739.3350, found 739.3352

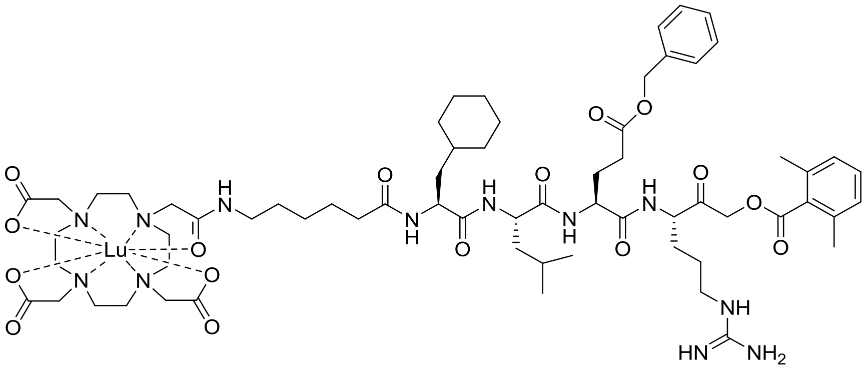

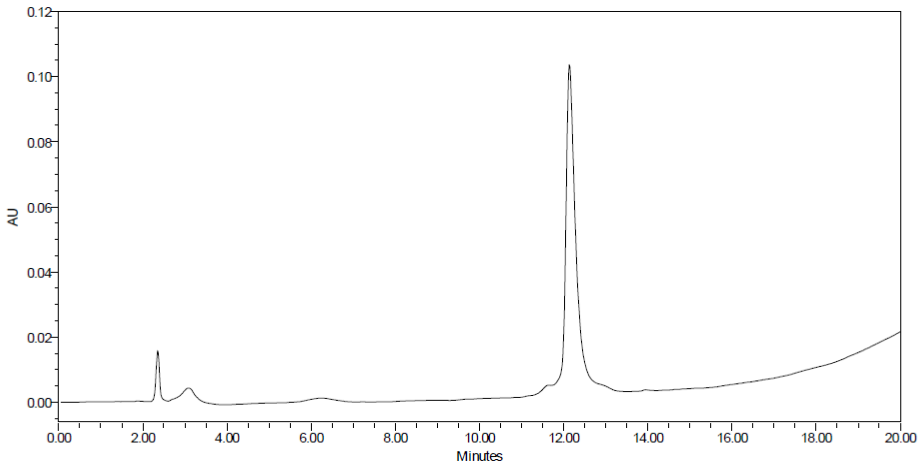

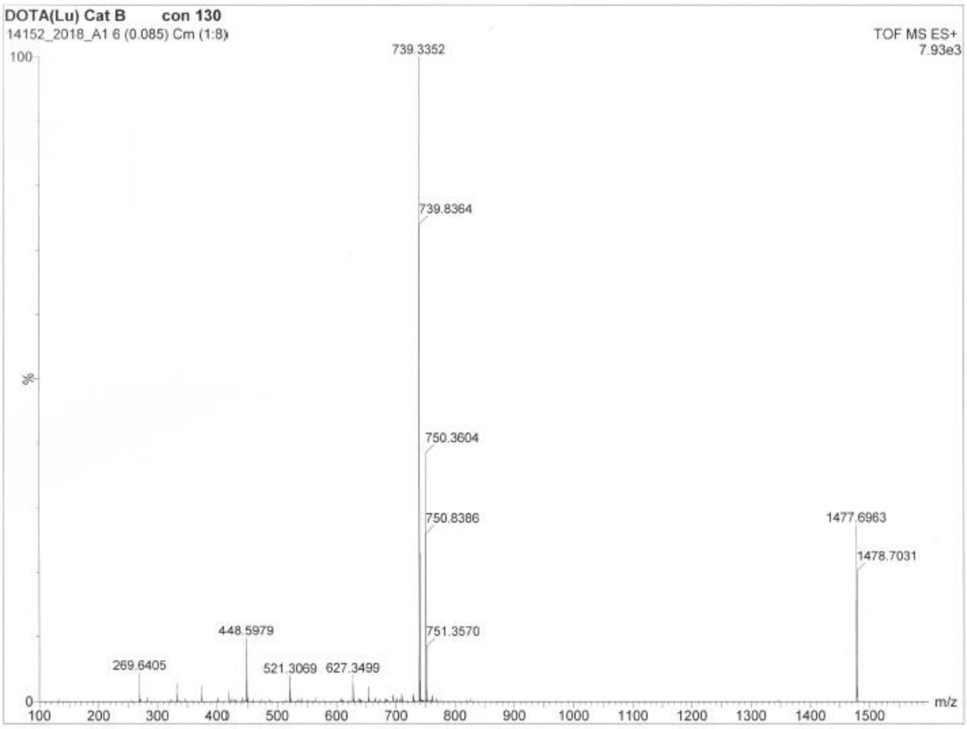


**DOTA(Gd)-Ahx-Asp-Glu-Val-Asp-AOMK**

HRMS (m/z): $[M+H]^+$ calcd for $C_{50}H_{72}GdN_9O_{20}$, 1277.4208, found 1277.5522

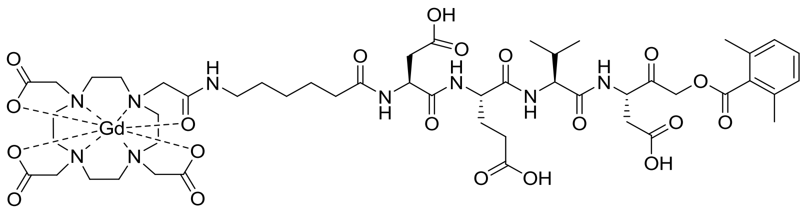

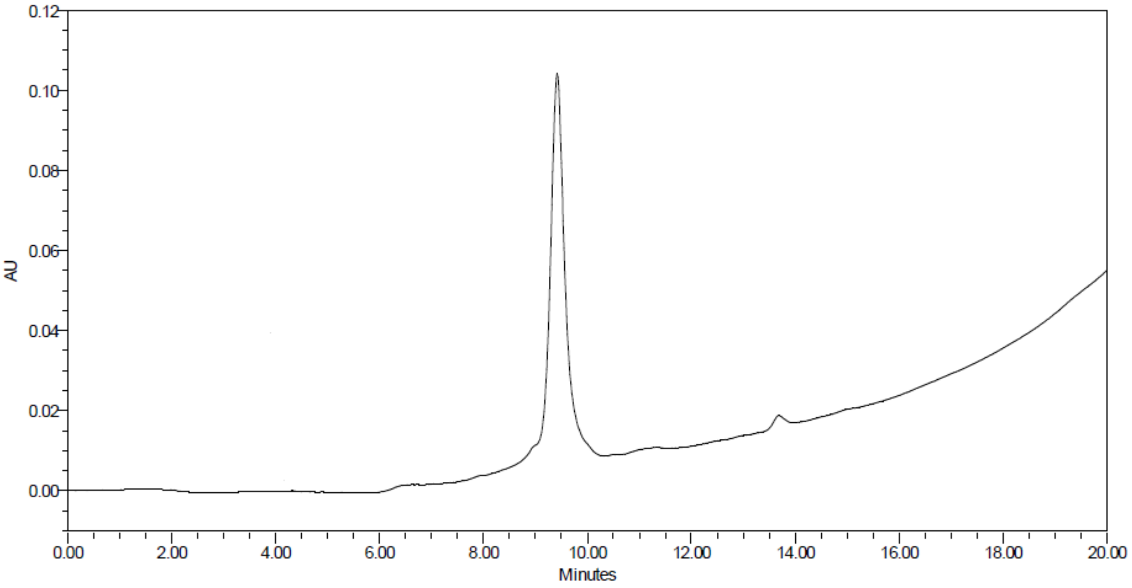

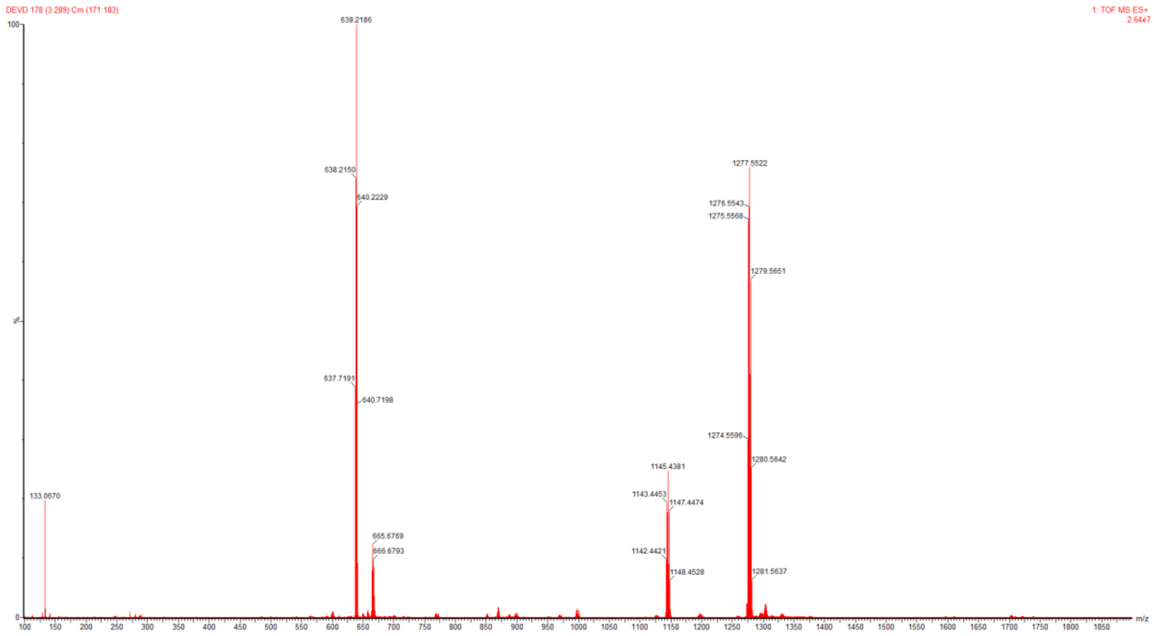
DEVD 178 (3.289) Cm (171:183)
1: TOF MS ES+
2.64e7
639.2186
638.2150
640.2229
637.7191
640.7198
133.0670
665.6769
666.6793
1277.5522
1276.5543
1275.5568
1279.5651
1274.5596
1280.5642
1145.4381
1143.4453
1147.4474
1142.4421
1148.4528
1281.5637
%
m/z
